# From proteins to species ranges: a framework for understanding thermal adaptation during range expansions

**DOI:** 10.1101/2025.09.09.675142

**Authors:** Saismit H. Naik, Emanuel A. Fronhofer

**Author notes:** Correspondence Details Emanuel A. Fronhofer, Institut des Sciences de l’Evolution de Montpellier, UMR5554, Université de Montpellier, CC065, Place E. Bataillon, 34095 Montpellier Cedex 5, France.

## Abstract

Species distributions are governed by both ecological and evolutionary processes. Traditionally, ecological factors have been the primary focus of species distribution studies, but recent work emphasizes the importance of rapid evolution through local adaptation. Here, we focus on adaptation to changing temperatures, which is one of the central challenges populations face today. Importantly, thermal adaptation may be affected by the underlying thermodynamics. Despite many existing models in the fields of thermal biology and spatial evolutionary ecology, there is little integrative theory. However, understanding and modelling the thermodynamic constraints on thermal adaptation is likely essential for more nuanced predictions of the impacts of climate change. By integrating molecular mechanisms and population dynamics in a unified modelling framework, we here study how temperature-dependent processes at the protein level influence the macroecological patterns of range expansions. Our results highlight the importance of the microscopic processes underlying thermal adaptation for capturing the evolutionary ecology of range expansions. Specifically, the molecular bases of thermal adaptation define how and how fast thermal performance can evolve, which determines range expansion speeds. In general, our framework predicts that adaptation to warmer temperatures will be easier than adaptation to cold. Our study underscores the necessity for more interdisciplinary work, combining molecular mechanisms with population dynamics in space in order to improve climate change modeling, enhance prediction accuracy and provide better information for management and conservation of natural populations.

**Significance Statement:** As global temperatures shift, species must adapt to new climates, but how molecular changes scale up to influence population- and ecosystem-level patterns is poorly understood. Here, we link mutations affecting protein stability and enzyme activity to species abilities to expand along temperature gradients. Our models show that adaptation is faster at warmer temperatures and more constrained in the cold, reflecting how mutations shape protein function. Thermodynamic effects amplify the impact of beneficial mutations at higher temperatures, potentially accelerating evolutionary responses. By connecting molecular biophysics to population dynamics and range expansion, this work provides a cross-scale framework for predicting how organisms respond to warming environments.

## Introduction

Temperature is a primary driver of biological processes (Brown et al., 2004). The hierarchical influence of temperature creates cascading effects across organizational levels: it determines protein reactions and metabolic rates, which impact birth, death, and feeding rates at the individual level. These temperature dependencies cascade up to determine population growth rates (Luhring and DeLong, 2017), densities, and further influence species interactions, and, ultimately, define geographic ranges across ecosystems (Knapp and Huang, 2022,; Fig. 1). As a consequence, increases in temperature due to global change are recognized as major drivers of species range shifts (Hickling et al., 2005).

**Figure 1:**
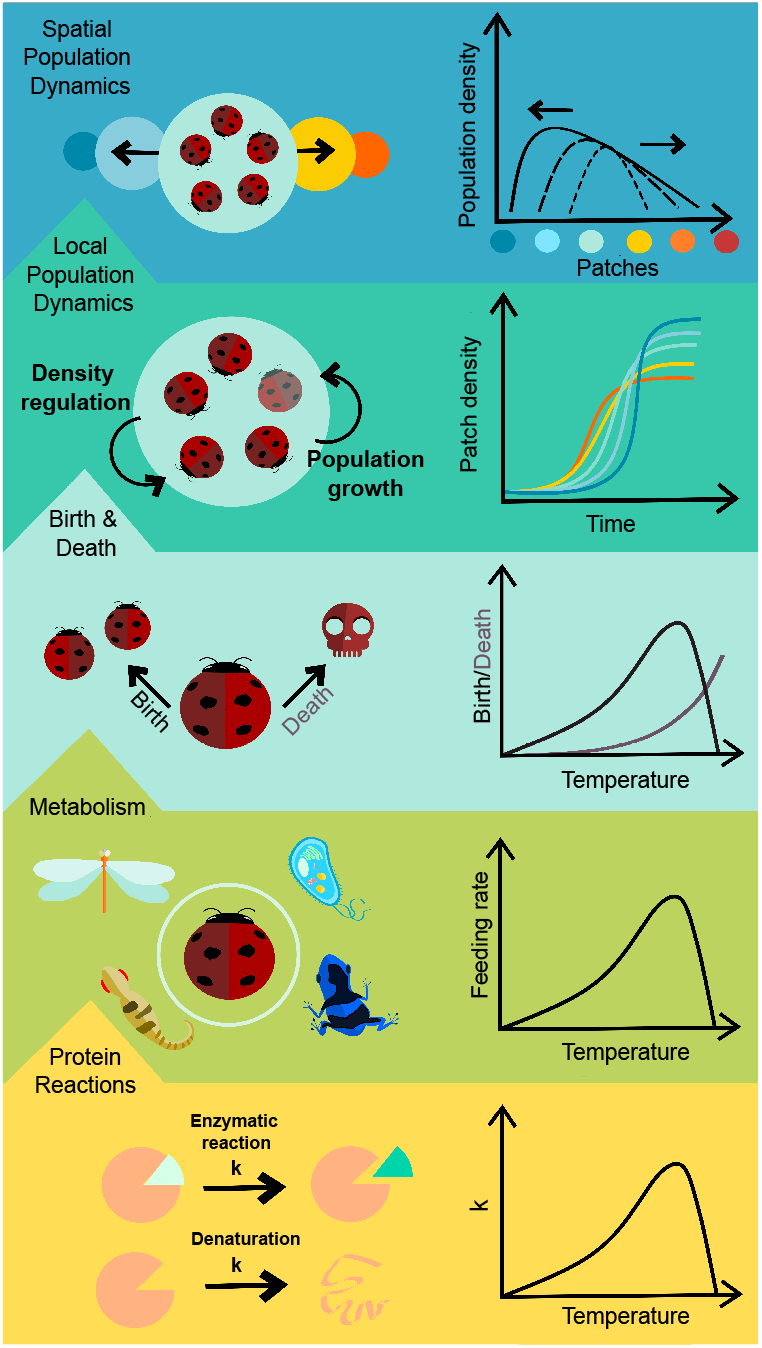
Metabolic rate is known to scale with temperature through the thermal dependence of its underlying protein reactions. Using well-established theories on the temperature-dependence of enzyme reactions, we build a framework that allows us to understand how scaling of protein reactions within their thermodynamic constraints propagates to population dynamics and even further to macroecological dynamics, including range expansions.

At an organismal level, thermal performance curves (TPCs) that describe how an organism’s performance changes with body temperature are the language of thermal biology. A thermodynamic framework has often been used to describe TPCs, including the popular metabolic theory of ecology (Brown et al., 2004) based on the exponential Arrhenius-Boltzmann model, which has been used to explain the temperature-dependence of many biological rates (Dell et al., 2011). Recent work by Arroyo et al. (2022) proposed a general theory of temperature-dependence in biology based on a model of entropy changes during protein reactions, fitting biological data across scales.

Surviving in changing temperatures, especially if these changes are large enough to prevent phenotypic plasticity (that is, the width of the TPC in this context) from being sufficient for buffering the change, hinges critically upon the biochemical properties of proteins and their ability to change evolutionarily to maintain functionality. Thermal adaptation becomes evident in comparisons across species, where numerous studies highlight striking patterns in thermodynamic properties. Essential metabolic enzymes (Rank and Dahlhoff, 2002; Niitepld, 2010; Meemongkolkiat et al., 2020) demonstrate temperature-associated clines in insect species, for example, where distinct gene variants balance enzyme efficiency and heat tolerance. In Antarctic and South American fish (Fields and Somero, 1998) flexibility near the active site in cold-adapted enzymes enhances catalytic efficiency by easing conformational shifts, which trades off with binding strength. Parallelism in patterns of protein-level adaptation across taxa (reviewed in Bomblies and Peichel, 2022) underscores the thermodynamics of protein structures as a powerful framework for understanding biochemical and biophysical adaptations and their influence on broader physiological and ecological processes.

Here, we aim at building a modelling framework that allows the integration of microscopic molecular evolutionary changes with macroscopic ecological dynamics in order to understand the consequences of thermal adaptation across scales. We adopt a pluralistic approach to modelling thermal dependence across biological scales that goes beyond Arroyo et al. (2022) by considering two major proposed models of growth rate TPCs that capture alternative biochemical mechanisms responsible for the shape of TPCs: First, the Protein denaturation-based model, taken from Chen and Shakhnovich (2009), suggests that the thermal evolution of individual growth rates occurs by modulating the copy numbers of folded proteins. This model links an organism’s replication rate to the functionality of each protein encoded by essential genes. It is supported by trends observed in viruses, bacteria, and insects (Buckley and Kingsolver, 2021; Frazier et al., 2006; Knies et al., 2009). Second, Macromolecular Rate Theory (MMRT) proposed by Hobbs et al. (2017) describes the unimodality of enzyme rate kinetics based on transition state theory (Eyring, 1935). Unlike the Protein denaturation-based model, it attributes the shape of the TPC to the temperature-dependence of the enthalpy and entropy changes accompanying enzyme-substrate reactions. This model is based on the observation that enzyme activity can decrease at temperatures below their denaturation temperatures, particularly in psychrophilic enzymes (Feller and Gerday, 2004). MMRT has been used to characterise soil microbial growth rates (Alster et al., 2016) and leaf respiration (Liang et al., 2018), for example.

Even though the above models and their extensions have been used to describe TPCs (Alster et al., 2016; Liang et al., 2018; Chen and Shakhnovich, 2010; Arroyo et al., 2022), the eco-evolutionary impacts of the underlying assumptions are currently not well understood. Hence, we apply scaling functions derived from the mechanistic models (MMRT and the Protein denaturation-based model) of protein thermal stability to a density-dependent population growth model. By introducing this mechanistic temperature-dependent population growth model into the center of a one-dimensional metapopulation model across a linear thermal gradient, we investigate the macroscopic consequences of assuming different TPC models for range expansion dynamics (Fig. 1) across both increasing and decreasing temperatures. To understand the contribution of both the skewed shape of the TPC as well the impact of assuming distinct processes underlying it, we consider a TPC defined by a Gaussian curve as our control (see details in Methods). Importantly, our framework is eco-evolutionary and allows for the evolution of thermal adaptation mechanisms. Using our pluralistic framework comparing the three models, we highlight the importance of asymmetry in TPCs and demonstrate the importance of considering emergent mutation sensitivity maps, resulting from the above-mentioned thermodynamic assumptions, to understand the effect of temperature on eco-evolutionary range dynamics.

## Results and discussion

### Thermodynamics set ecological range limits

Regardless of the TPC model, populations lacking thermal adaptation are unable to occupy the full environmental temperature gradient (Fig. 2 A), as their initial TPCs do not span the entire temperature range (Fig. S2 A). Importantly, the direction of range limitation, whether towards hotter or colder environments, depends on the specific biophysical mechanism: In the Protein denaturation-based model, the Gibbs free energy of protein unfolding 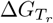 scales with temperature (Eq. S8) and must remain negative to represent a spontaneous reaction. This results in a sharp boundary along the hot front. As a consequence, the Protein denaturation-based model restricts expansions into hotter environments.Similarly, in the Enzyme catalysis-based model, the entropy change parameter 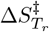 also scales with temperature (Eq. S6) and must remain negative. However, the consequence of this constraint is different: The Enzyme catalysis-based model constrains expansion into colder environments (Fig. 2 A).

**Figure 2:**
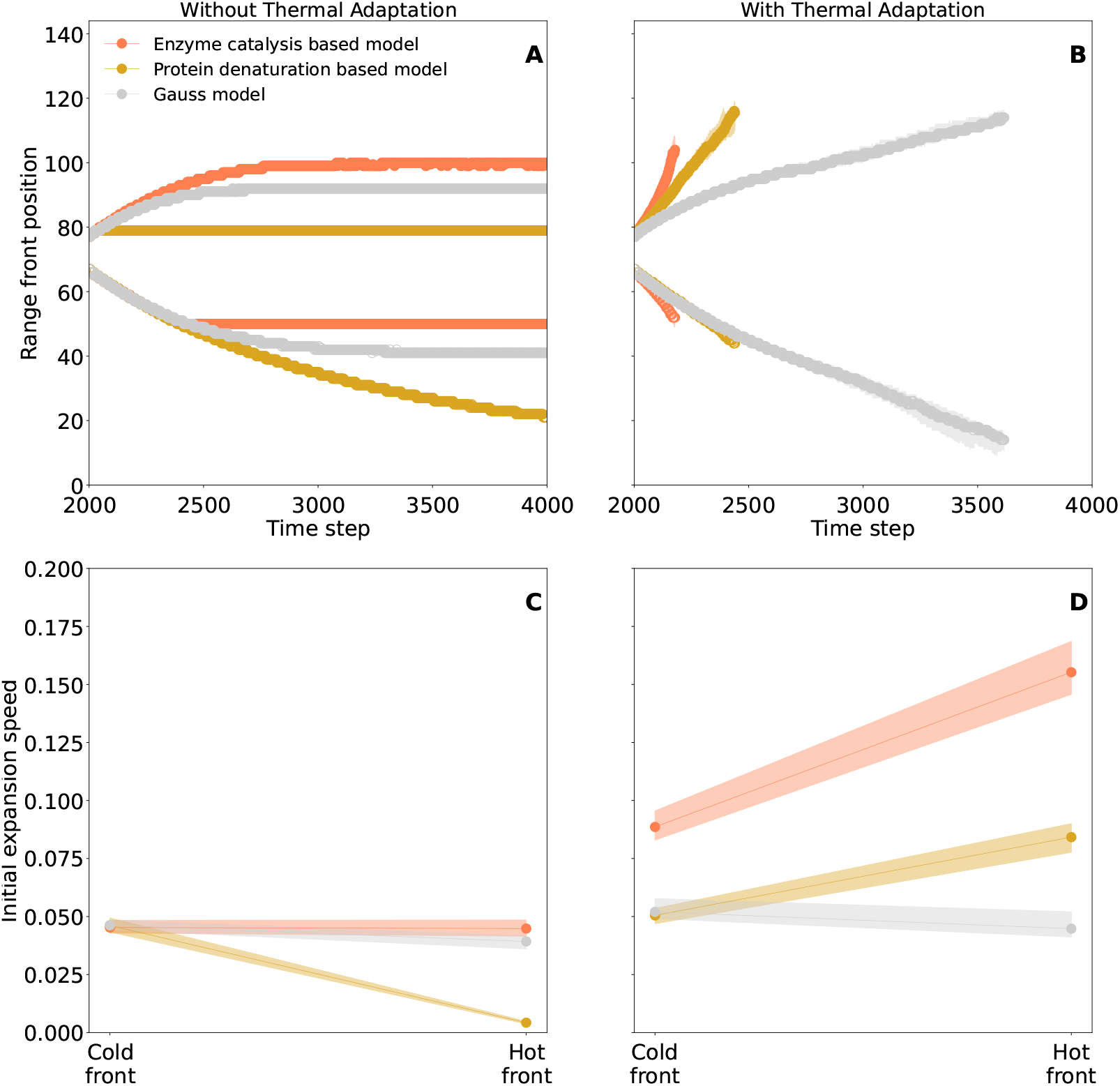
Range expansions without (A, C) and with thermal adaptation (B, D). Without thermal adaptation the different models reach different range limits (A) highlighting model-specifc constraints. Expansion speeds (C) do not differ markedly between hot and cold front except for the Protein denaturation-based model. With thermal adaptation expansions are faster and populations spread further into the gradient (B). The thermodynamically motivated models show faster expansions towards the hot side (D). Range front positions (A, B) are averaged across 80 replicates. The last 3 occupied patches with more than 10 individuals are considered the range front. Initial speeds of expansion (C, D) are calculated by fitting a linear regression to patch front dynamics before 2100 time steps. We show medians and interquartile ranges (shaded region) among replicates. Population growth parameter values can be found in Table S1. Temperature scaling parameters can be found in Table S2.

### Evolutionary constraints determine range expansion speeds

With thermal adaptation, thermodynamic constraints can be overcome in both thermodynamic models. With evolution, both thermodynamically motivated models show faster hot than cold range expansion. The Gaussian model, which we here include as a control and an often used non-mechanistic alternative (Angilletta Jr, 2006; Asbury and Angilletta Jr, 2010), continues to show symmetrical range expansion (Fig. 2 B, D).

Two distinct mechanisms underlie these trends: First, adaptation at the cold front is more constrained in the thermodynamic models. In Fig. 3 A, B we observe minimal cold adaptation in median TPCs at range ends in the thermodynamic models, while the non-mechanistic Gaussian model in Fig. 3 C is able to shift its thermal performance curve toward cooler temperatures. We investigate the thermal trait dynamics underlying niche evolution in more detail in the next section.

**Figure 3:**
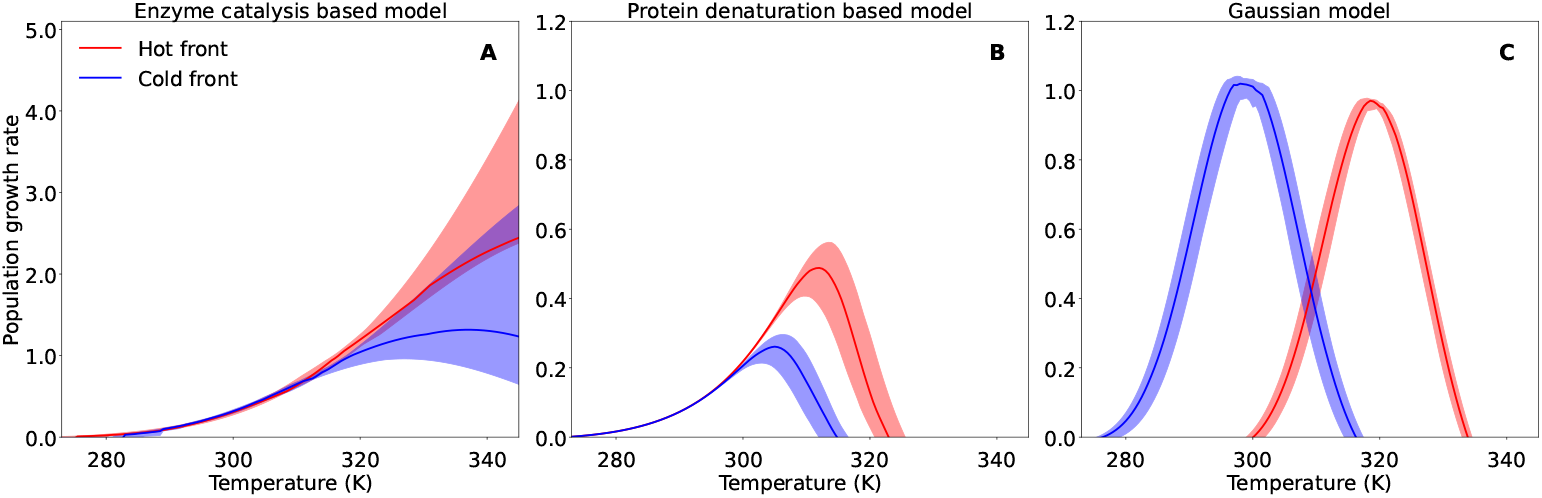
Evolved TPCs at the range fronts (hot side: red; cold side: blue). The two thermodynamically motivated models (A: Enzyme catalysis-absed model; B: Protein denaturation-based model) show asymmetric and hotter-is-better trends, while the Gaussian control (C) shows symmetrical adaptation to both hot and cold fronts. The lines show medians and the shaded area is the variance of the thermal niche across the 80 replicates. Population growth parameter values can be found in Table S1. Temperature scaling parameters can be found in Table S2.

Second, thermodynamic models show a clear hotter-is-better trend at the hot front (Knies et al., 2009). Hotter-is-better is a hypothesis of thermal adaptation that populations adapted to warmer temperatures will have higher maximum growth rates than those adapted to low temperatures. Thermodynamic rate depressing effects of low temperatures cannot be compensated by plasticity or genetic changes. In our simulations, populations following the thermodynamic models evolve higher maximum growth rates as they adapt to warmer conditions (Fig. 3 A, B). To understand these asymmetries in niche evolution, we next compare the dynamics of the underlying thermal traits.

#### Asymmetric mutation sensitivity

To better understand why cold-end adaptation is limited compared to hot-end adaptation in thermodynamic models, we examined the distribution and effect sizes of mutations across the landscape (Fig. 4). Thermodynamic models show increasing absolute magnitudes of fitness effects of mutations with temperature as seen in Fig. 4 A and B. Inherently, this observation is not surprising as the mechanistic models show greater asymmetry in their TPCs than the Gaussian model: Steeper slopes when the thermal niche dips at hotter temperatures will automatically lead to bigger mutation effects for hotter temperature than for lower temperatures for the same shift in the curve.

**Figure 4:**
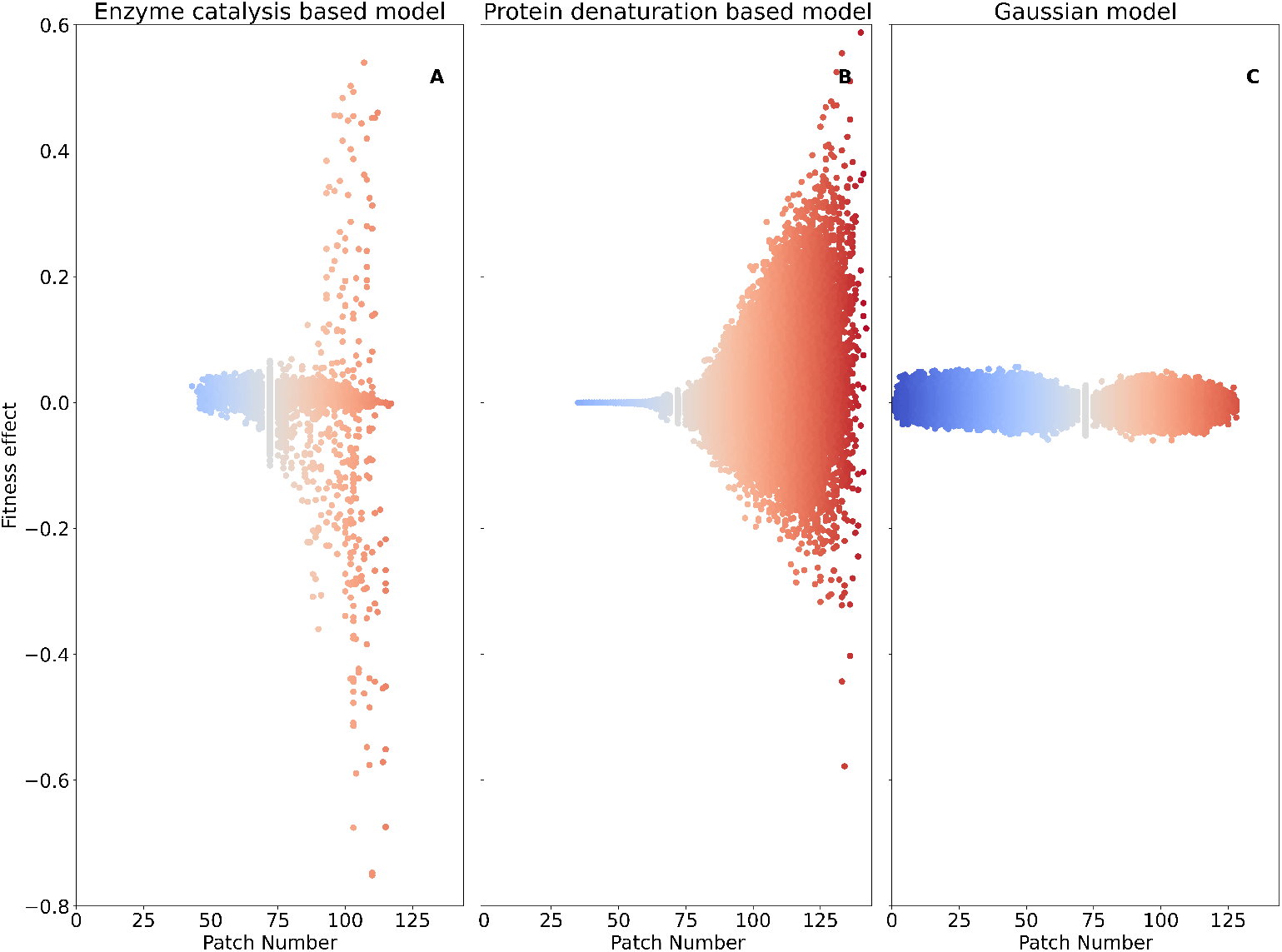
Spatial distribution of fitness effects. By tracking every mutation that occurs in our simulation, we are able to visualise the distribution of associated fitness effect over space. The color of the points corresponds to the patch temperature and is on a gradient from cool (blue) to warm (red). A) Enzyme catalysis-based model; B) Protein denaturation-based model; C) Gaussian model. Population growth parameter values can be found in Table S1. Temperature scaling parameters can be found in Table S2.

We do see differences between the two thermodynamic models, as the Enzyme catalysis-based model shows a wider distribution of mutation effects with hotter temperatures (Fig. 4 A, B). However, it is important to note that these specific differences depend on parameter values: Fig. S11 explores this in detail and shows how mutation effects dependent both on the mean of the trait around which the mutation occurs as well as on the impact of temperature. The Protein denaturation-based model operates in a less sensitive area (shaded green region) of parameter space in Fig. 2 (Fig. S11 C, D), limiting its adaptive response. This difference explains why the Enzyme catalysis-based model achieves higher overall expansion speeds once evolution is enabled (Fig. 2 D). In comparison, as expected, the Gaussian model shows symmetrical mutation effects in both Fig. 4 C and Fig. S11 E, F. As the mutation effects are dependent on individual thermal trait changes, we plot mutations that result due to changes in single thermal traits, and we show the contribution of individual traits to the fitness effects in Fig. S12, S13 and S14. Following Fig. S11, we confirm a higher sensitivity to mutation with temperature in both thermodynamic models (Fig. S12, S13), while the Gaussian model (Fig. S14) has symmetric mutation effects. We also note that in each model, certain traits may have higher magnitudes of mutation effects compared to the others.

Despite these specificity, our analysis allows us to conclude that faster range expansions to the hot side are a general feature of our thermodynamically motivated models. Quantitative differences, as seen in the mutational effects (Fig. S11 A, B), for example, will depend on the choice of parameters. Here, we start our simulations with TPCs that have similar median behavior and mutation effects in order to make comparisons easier.

#### Predictability

Apart from the speed of range expansion, evolution can also impact variability between replicates during range expansion (Williams et al., 2019a). To characterize the predictability of expansion dynamics, we examined the variability in range front trajectories across replicate simulations. In Fig. S10 A, thermodynamic models show greater trajectory dispersion along the hot side, with a corresponding increase in interquartile range over time (Fig. S10 B). We find that the Enzyme catalysis-based model shows most variability, along the hotter side followed by the Protein denaturation-based model and the Gaussian model. These results are concordant with the greater distribution of fitness effect sizes seen in the Enzyme catalysis-based model in Fig. 4 A) which amplify stochastic differences between individual replicates, leading to less predictable dynamics.

### Dispersal evolution

As shown previously, local adaptation is not the only trait that may evolve during range expansions. The second critical trait is dispersal. Therefore, in order to understand the robustness of the results presented above and the impact of both dispersal evolution and thermal adaptation occurring simultaneously, we alos allowed dispersal to evolve. Importantly, we do not see an overall change in the trends in range front dynamics as observed above (Fig. S15). Notably, the time to reach the end of landscapes is not markedly different despite evolving dispersal probabilities. Finally, we consider the niches at hot and cold fronts at the end of the simulations (Fig. S16 as compared to Fig. 3). These results indicate that dispersal evolution is secondary to thermal adaptation during range expansions along a thermal gradient, as has been shown before by Deshpande and Fronhofer (2022), for example. For a more detailed discussion of dispersal evolution and its relevance for biodiversity maintenance during global change see Kamal et al. (2025).

## General discussion

Our study takes a multi-scale approach to modeling evolution of thermal performance curves and demonstrates how biophysical constraints to thermal adaptation of crucial protein level properties can propagate up to range expansions along a thermal gradient (Fig. 1). Specifically, we develop models of range dynamics that account for temperature dependence of processes critical to survival under changing thermal regimes: First, protein stability (Eq. S7) and second, enzyme catalysis rates (Eq. S22). By analyzing these models, we find that larger mutation effects in hotter temperatures predicted by first-principles thermodynamic models lead to faster expansion along increasing temperatures and limit adaptation along decreasing temperatures. Our results highlight how constraints and characteristics of protein-level adaptations cascade all the way to macroecological patterns of range dynamics (Fig. S16). Below we discuss our results within the cross-scale perspective presented in Fig. 1.

### Many roads lead to Rome: Thermal and phylogenetic context of mutation effects

By mapping scaling functions from thermodynamic models to growth rate scaling of populations, our framework allows us to link thermodynamic protein traits both to fitness and to how mutations in these traits change fitness across temperatures. This mapping (Fig. S11) highlights how temperature and mean trait values interact to shape mutation effects.

Empirical studies have found that the fitness effects of mutations are not fixed but instead depend on both thermal context and evolutionary history. For example, several studies report stronger mutation effects at higher temperatures (Berger et al., 2021; Chen and Zhang, 2023; Chu et al., 2020), in line with the Stronger Beneficial Mutation Effects Hypothesis (Chen and Zhang, 2023). Recent empirical studies have found that the fitness effects of mutations depend on the environmental temperature. For example, several studies report stronger mutation effects at higher temperatures (Berger et al., 2021; Chen and Zhang, 2023), in line with the Stronger Beneficial Mutation Effects Hypothesis (Chen and Zhang, 2023). This hypothesis suggests that hotter temperatures amplify the fitness impacts of beneficial mutations without reducing fitness at colder temperatures, producing the Hotter is better pattern commonly discussed in thermal adaptation studies (Malusare et al., 2022).

Recent evidence for this hypothesis comes from Chen and Zhang (2023), who showed that *E. coli* exhibited stronger beneficial effects at warmer assay temperatures. However, the same experiment also revealed an important nuance: in *P. fluorescens*, adaptation to high temperatures was associated with trade-offs in cold performance, highlighting that multiple mechanisms of thermal adaptation can occur in parallel (Razvi and Scholtz, 2006).

Our results mirror this diversity. Depending on the starting position of thermal phenotypes, thermodynamic models predict both scenarios: mutations that amplify fitness at high temperatures (seen in Fig. S11 A, B and E) and mutations that generate trade-offs at low temperatures (or vice versa; Fig. S11 D, also see fitness effects along cold gradients in Fig. 4 A versus B). Another example is reported by Nguyen et al. (2017), who explore how enzymes adapted over evolutionary time to maintain catalytic efficiency as Earth’s temperatures cooled dramatically. The study focused on adenylate kinase (Adk), an essential enzyme, using ancestral sequence reconstruction to study three billion years of evolution from ancient thermophilic ancestors to modern mesophilic and psychrophilic variants. The authors show that adaptations altering heat capacity changes in the Adk enzymes arose gradually and specifically in certain lineages. These adaptations were contingent on previously accumulated mutations that influenced protein folding dynamics and the enzyme’s internal network of stabilizing interactions such as salt bridges.

Our results show that, even without explicit epistasis, the thermodynamic models can have a diversity of outcomes (Fig. S11). In this sense, our framework provides a mechanistic explanation for the range of empirical patterns observed across taxa and evolutionary histories. An exciting future step in theoretical studies on thermal adaptation to new environments would therefore be to include context-dependent thermodynamic constraints that may be derived from both history of exposure to stressful environments and phylogeny (Cui et al., 2025; Zhang et al., 2025; Kontopoulos et al., 2020). Microbial systems are emerging as an excellent system to study the impact of thermodynamic constraints on thermal adaptations, especially combined with methods like Flux Balance Analysis and Genome Scale Metabolic Models that can give more detailed information on metabolic networks underlying responses to thermal stress (Alster et al., 2023; Qiu et al., 2023).

### The road from Rome: Thermal phenotypes and population-level consequences

Thermal dependence of rate limiting metabolic processes can map to growth and survival, and mutations affecting these processes alter growth and survival in distinct ways. These vital rates ultimately shape population dynamics in space and time. Thermal ecology has long studied how temperature-dependence in growth and survival influences population-level outcomes, but explicitly incorporating underlying molecular mechanisms or their evolutionary change is an ongoing challenge (Bernhardt et al., 2018; Usui and Angert, 2024a).

By adopting an explicitly molecular perspective, our model demonstrates how range dynamics result from the interaction of thermodynamic constraints, mutation effects, and evolutionary responses. In particular, we show that increasing mutation effect sizes at higher temperatures increases variability in growth (Fig. 4), reduces equilibrium densities (Fig. S2), and accelerates turnover. These predictions are partly consistent with empirical results such as the range expansion of duckweed along a thermal gradient reported by Usui and Angert (2024b), which showed increased variability at warmer conditions. However, our simulations do not reproduce the faster spread and higher densities at the range front seen in the experiment, highlighting how assumptions about the scaling of competition and density dependence (e.g., competition scaling with birth rates; Eq. 7) critically shape outcomes. While such assumptions have some empirical support (Fronhofer et al., 2024; Johnson et al., 2016), a more integrative understanding of how birth, death, and competition scale with temperature is needed (Amarasekare and Coutinho, 2014; Stockseth et al., 2025).

An important consequence of this interplay is the predictability of range expansion dynamics. As reported here, evolution can decrease the predictability of range expansion, depending on the balance between variance-reducing processes (e.g., natural selection) and variance-increasing processes (e.g., gene surfing) (Williams et al., 2019b). According to the framework proposed by Williams et al. (Williams et al., 2019b), density at the expanding range front plays a critical role for predictability: shallow fronts foster variation-generating evolution, while steep fronts promote variation-reducing evolution.

### Rome in the eye of the beholder: Cross-taxa trends on thermal evolution

Our results highlight the need for a comparative perspective on thermal adaptation across taxa. While thermodynamic models capture key microbial dynamics, where single rate-limiting processes or mutations in thermally sensitive proteins can directly shift thermal tolerance (Chen and Shakhnovich, 2010; Alster et al., 2018, 2016; Leuenberger et al., 2017), range expansion dynamics in higher taxa appear to follow different rules. In microorganisms, single-locus changes such as those affecting protein stability or chaperone function can measurably shift upper thermal limits (AlZaben et al., 2021; Lehmann et al., 2023; Madern et al., 2024; Pinney et al., 2021). By contrast, in larger organisms, evolutionary responses are slower, more strongly shaped by standing genetic variation, and more dependent on evolutionary history and past temperature regimes. This difference likely reflects a time-scale separation between ecology and evolution, as well as the polygenic nature of thermal adaptation in multicellular systems.

Our model predicts faster expansion along warming gradients under assumptions of protein-property adaptation and feeding-rate scaling. Yet these outcomes diverge from macroscopic empirical patterns in complex taxa, where adaptation to hotter environments is generally more constrained than adaptation to colder ones (van Heerwaarden et al., 2016; Campbell-Staton et al., 2017; Overgaard et al., 2011). Across latitudes, minimum tolerable temperatures (*CT*_*min*_) show greater variability and acclimation capacity than maximum tolerable temperatures (*CT*_*max*_) (Arajo et al., 2013; Qu and Wiens, 2020). This asymmetry has been explained by stronger physiological constraints at the upper thermal limit, as well as by the fact that most species occupy climatic niches below their *CT*_*max*_, thereby experiencing greater variation at lower temperatures (Qu and Wiens, 2020). Further work is needed to clarify the relationship between physiological tolerance and realized climatic niches, as highlighted in Qu and Wiens (2020) as well as the importance of microclimatic factors. These asymmetries may also underscore how adaptation to cold and hot environments may be governed by different processes (Tattersall et al., 2012).

The enzyme cytosolic malate dehydrogenase (cMDH) illustrates these parallels and divergences. Adaptive changes in cMDH have been observed in both microorganisms (Madern et al., 2024; Hobbs et al., 2017) and higher organisms (Zhang et al., 2025). Yet, while microbial adaptation often follows directly from molecular changes, larger organisms experience additional layers of complexity due to polygenic adaptation, pleiotropic trade-offs, and phylogenetic constraints (Zhang et al., 2025; Bomblies and Peichel, 2022).

In this sense, while the roads to Rome may begin with conserved molecular mechanisms, the routes taken by microbes and higher taxa diverge in how these mechanisms translate to population growth, range expansion, and macroecological patterns. Future work that integrates molecular constraints with polygenic processes and evolutionary history will be crucial for explaining why thermal adaptation manifests so differently across the tree of life.

## Conclusion

Our work, by singling out protein-level adaptation, bridges biophysical adaption to ecology and shows that simple mechanisms can capture many trends observed in thermal adaptation studies. Our results reveal that thermodynamic constraints on metabolic processes create asymmetric range limits and speeds of range expansion along temperature gradients, favoring adaptation and faster spread at higher temperatures. By using mechanistic models, we show that both mutation effects and population dynamics fundamentally shape not only the rate but also the variability of adaptation. These insights underscore the need for detailed, mechanism-based approaches to predict evolutionary and ecological responses under climate change combining studies on biophysical adaptations to novel environments to population growth and finally to adaptation during range expansions.

## Modelling framework

We develop a continuous-time individual-based model to simulate the population dynamics of an asexually reproducing species expanding its range along a temperature gradient in a patchy landscape (stepping stone metapopulation model).

### Temperature dependence of demography

#### Introducing the temperature scaling function (*s*(*p*_*evo*_, *T*))

The continuous-time Beverton-Holt model can be derived by assuming that resources quickly equilibrate in a chemostat model (Thieme, 2003). The dynamics of the focal consumer population are then as follows:

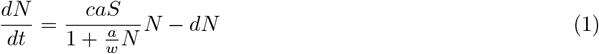

where *w* is the flow rate of resources in and out of the system, *S* is the constant resource flowing into the system, *c* is the efficiency of resource consumption, *a* is the feeding rate. Setting *b* = *eaS* and 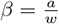, we get:

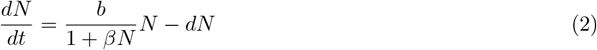

To understand how the system will depend on temperature, we consider the feeding rate *a* to be the temperature-dependent parameter, as noted in the literature (del Giorgio and Cole, 1998). As both *b* and *β* are constant multiples of *a*, they scale identically with temperature in our model. The temperature scaling function *s*(*p*_*evo*_, *T*) takes as arguments the local temperature *T* and its evolving parameters *p*_*evo*_. In general, the temperature-dependence of birth rate and intraspecific competition coefficient is given by:

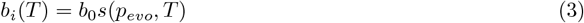

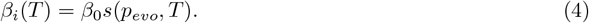

The function *s* is given by the Protein denaturation-based model (Eqn. S7), the Enzyme catalysis-based model (Eqn. S22), and a Gaussian curve as a non-mechanistic reference (Eqn. S23) as described in detail in the Supplementary Information. For biological relevance, we have used values relevant to microbial communities. The values *b*_0_ and *β*_0_ are taken from Fronhofer et al. (2024), who used data from *Tetrahymena thermophila* at 20°C and can be found in the Table S1.

#### Evolving parameters (*p*_*evo*_)

For each model, we consider parameters that are central to the underlying thermodynamic mechanism and for which evolutionary adaptation has been empirically observed. In the Protein denaturation-based model, these include the Gibbs free energy of unfolding (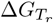) and the entropy of unfolding (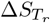), which together determine the thermal stability of essential proteins. In the Enzyme catalysis-based model, the key evolving parameters are the activation enthalpy (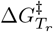), activation entropy (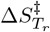), and the activation heat capacity (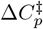), which influence catalytic efficiency across temperatures. Adaptation in these parameters has been documented in empirical systems. These changes reflect selective pressures that optimize molecular function under thermal constraints. Further discussion and supporting literature are provided in Section of the Supplement.

In the simulations with evolution, the evolving parameters are initialised with Gaussian standing genetic variation in the evolving parameters. The details of the distribution for each model are given in Table S2. At each birth, the traits may mutate with a probability 0.05. In case of a mutation, the new trait values are chosen from a Gaussian distribution whose mean is the same as the mean of the parent individual, and the standard deviation of each model is given in Table S2.

We also ensure that the effects of mutations are comparable between models. The mutation standard deviations for each model are adjusted so that the interquartile range of the mutation effects, derived from sampling model parameters, remains consistent across all thermodynamic models. This alignment ensures that differences in mutation impact are not driven by discrepancies in model parameterisation.

#### Introducing the temperature scaling of death rate

Temperature dependence of the death rate is taken to be of the Arrhenius-Boltzmann equation form, as has been widely observed (McCoy and Gillooly, 2008):

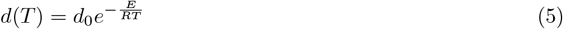

#### A temperature-dependent demographic model

Hence, our temperature-dependent population growth model is:

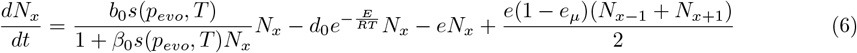

The equilibrium density then is given by:

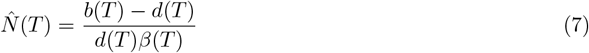

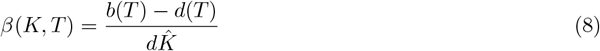

Considering the temperature dependence of the parameters, we obtain the following expression for equilibrium densities:

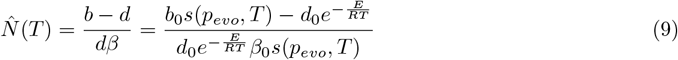

Note that Eqn. 9 only holds without dispersal.

### Range expansion model

As described above, our model is based on a continuous-time Schoener population growth model (Fronhofer et al., 2024) with emigration assuming a stepping stone metapopulation model. In the absence of trait evolution, the mean expectation of the individual-based model is the same as the dynamics of the following equation:

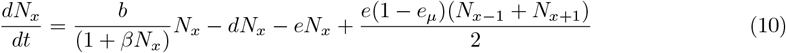

where *N*_*x*_ is the population size of the species in the *x*^*th*^ patch, *b* is the species’ birth rate, *d* is the death rate, *e* is the emigration rate, *e*_*µ*_ is the emigration mortality (Bonte et al., 2011) and *β* is the intraspecific competition coefficient.

### Initialisation and analysis

We used the modified Gillespie algorithm by Allen and Dytham (2009) to simulate the above equations as an individual-based model (see Supplementary Informationfor details on the implementation).

#### Landscape

For simplicity, we consider a linear landscape with 144 connected patches. Patch temperatures start from 0 °C and increase with a 0.5 °C step, till 72 °C.

#### Controls

To keep relevant controls, we implement a full factorial modelling design crossing thermal adaptation (present and absent) with dispersal evolution (present and absent) for all three TPC models (Protein denaturation-based model (Eqn. S7), Enzyme catalysis-based model (Eqn. S22) and the Gaussian Curve (Eqn. S23).

#### Initialisation

We consider the Gaussian model to have an optimum at 36°C (approximate optimum temperature for *Tetrahymena* (Prescott, 1957)) with an approximate range between 20°C and 45°C (Weber de Melo et al., 2020).

The Protein denaturation and Enzyme catalysis-based models are parameterized using the Gaussian model as a reference between the temperatures 20°C and 45°C. The `differential_evolution optimization` algorithm from the SciPy library was used to fit these models optimally. By fitting both models to the Gaussian curve, we reduce variability introduced by differences in initialization. The resulting curve can be seen in Fig. S2.

The metapopulation model consists of a 144-patch linear landscape. The central 10 patches, that is, the patches with temperatures from 33.5 °C to 38.5 °C are initialised with 200 individuals each. Dispersal to either end of the landscapes is not allowed for the first 2000 time steps to let the individual traits evolve to an equilibrium in the central patches. After 2000 time steps of burn-in, which corresponds approximately to 9500 generations on average in the ecological control with our baseline parameterised at 20 °C, individuals can disperse across the whole landscape.

#### Replicates and stopping simulations

All models were run for the three scenarios with 80 replicates for each and the temperature scaling model described in the previous section. The simulations are stopped when the first replicate reaches either end of the landscape in order to focus on the range expansion phase only.

## Author contributions

EAF and SHN conceived the study. SHN analysed the mathematical models. SHN and EAF wrote the manuscript.

## Acknowledgements

This work was supported by a grant from the Agence Nationale de la Recherche (No.: ANR-19-CE02-0015) to EAF. This is publication ISEM-YYYY-XXX of the Institut des Sciences de l’Evolution Montpellier.

## Declaration of AI use

We declare having used ChatGPT 4.0 for editing and grammar checks.

## Data availability

Code will be made available via GitHub and a Zenodo DOI.

## Notes

### Competing Interest Statement

The authors have declared no competing interest.

