## Supplementary Materials for "From proteins to species ranges: a framework for understanding thermal adaptation during range expansions"

#### Supplementary Theory

##### Modelling the temperature scaling function ($s(p_{evo}, T)$)

In 1889, Svante Arrhenius introduced a model describing the exponential temperature dependence of chemical reaction rate constants (Arrhenius, 1889) , which has been used widely to express thermal dependence of biological rates, from chemical reactions to growth rates (Dell et al., 2011; Gillooly et al., 2001). The equation is expressed as :

$$k_r = Ae^{-\frac{E}{k_B T}} \tag{S1}$$

where $k_r$ the reaction rate constant $A$ represents the frequency of reactant collisions, $E$ is the activation energy, $k_B$ is the Boltzmann constant, and $T$ is the temperature. The Arrhenius model is mechanistic in its use of the Boltzmann factor but lacks a theory for the activation energy $E$, making it an empirical model unable to account for the unimodality of thermal performance curves (TPCs) seen in biological systems. The Eyring-Evans-Polanyi (EEP) (Eyring, 1935) transition state theory (TST) later provided a more comprehensive and mechanistic framework, especially for enzyme-catalyzed reactions, describing them through:

$$k_r = \frac{k_B T}{h} e^{\frac{-\Delta G^{\ddagger}}{RT}} \tag{S2}$$

where $k_r$ denotes the reaction rate constant, $k_B$ is the Boltzmann constant, $h$ is Planck's constant, $\Delta G^{\ddagger}$ is the Gibbs free energy of activation, and $R$ is the universal gas constant. The term $\frac{k_B T}{h}$ is a pre-exponential factor representing the number of successful collisions per second that reach the transition state. The change in Gibbs free energy of activation ($\Delta G^{\ddagger}$) is related to two key thermodynamic parameters: 1) Activation enthalpy ($\Delta H^{\ddagger}$): Represents the energy needed to reach the transition state, analogous to the activation energy in the Arrhenius model (Eqn. S1). 2) Activation entropy ($\Delta S^{\ddagger}$): Represents the change in the disorder or randomness when moving from reactants to the transition state. Their relationship is expressed as:

$$\Delta G^{\ddagger} = \Delta H^{\ddagger} - T\Delta S^{\ddagger}$$

A positive $\Delta S^{\ddagger}$ value suggests an increase in disorder at the transition state, which generally enhances the reaction rate, while a high $\Delta H^{\ddagger}$ value indicates a higher energy barrier, slowing the reaction. The enthalpy ($\Delta H$) and entropy ($\Delta S$) of reactions are often temperature-dependent, which in turn affects population growth rates (Arcus et al., 2016). A more accurate representation of this temperature-dependence involves considering a constant heat capacity change $\Delta C_p$, which defines temperature scaling $\Delta H$ and $\Delta S$ of reactions, leading to:

$$\frac{d(\Delta H^{\ddagger})}{dT} = \Delta C_p^{\ddagger} \tag{S3}$$

$$T\frac{d(\Delta S^{\ddagger})}{dT} = \Delta C_p^{\ddagger} \tag{S4}$$

Therefore $\Delta H(T)$ and $\Delta S(T)$define enthalpy and entropy changes at temperature $T$ are given as:

$$\Delta H(T)^{\ddagger} = \Delta H_{T_r}^{\ddagger} + \Delta C_p^{\ddagger}(T - T_r) \tag{S5}$$

$$\Delta S(T)^{\ddagger} = \Delta S_{T_r}^{\ddagger} + \Delta C_p^{\ddagger} \ln\left(\frac{T}{T_r}\right) \tag{S6}$$

with $T_r$ being the reference temperature. These equations are crucial for modeling how metabolic processes, and thus population growth rates, vary with temperature.

The EEP equations (Eqn. S2) have been extended to derive many unimodal models of population growth TPCs, such as done by Johnson and Lewin (1946a) and Schoolfield et al. (1981). Most recently, in a bid to provide a simple mechanistic model of the general temperature dependence of biological rates, Arroyo et al. (2022) considered the EEP equations and assumed unimodality of TPCs comes from the temperature dependence of $\Delta S_T^{\ddagger}$, that is, the conformational entropy of enzyme-substrate reactions. The assumption reduced the EEP equations to an exponential function modified by a power law and the authors were able to find good fits of TPCs across the biological organisational hierarchies. But assuming a single temperature-dependent quantity to describe the unimodality of TPCs falls short as we move from an ecological to an evolutionary perspective.

While simple mechanistic models, such as those proposed by Arroyo et al. (2022), can provide good fits to TPCs, they may not fully capture the complexities of thermal adaptation processes. Assuming that a single factor drives unimodality, potentially overlooks the nuances of how different biochemical properties adapt evolutionarily to temperature changes. To demonstrate this, we explore two distinct models based on Eqns. S2 that both explain TPC unimodality but yield different macroecological outcomes as they consider different biochemical properties of a single reaction/protein under selection: 1) thermal stability of a single protein and 2) temperature dependence of a single enzyme catalysis reaction.

#### Protein denaturation-based model

The Protein denaturation-based model proposed by Chen and Shakhnovich (2010) suggests that the population growth rate is directly proportional to the fraction of proteins in their native state. In this model, the Gibbs free energy of protein unfolding at temperature $T$ is denoted by $\Delta G(T)$, and the exponential scaling factor for growth rate is characterized by the activation enthalpy $\Delta H^{\ddagger}$:

$$s(p_{\text{evo}} = [\Delta G_{T_r}, \Delta S_{T_r}], T) = \begin{cases} b_0 \dfrac{e^{-\Delta H^{\ddagger}/RT}}{1 + e^{-\Delta G(T)/RT}}, & \text{if } \Delta G(T) < 0 \\ 0, & \text{otherwise} \end{cases} \tag{S7}$$

where $b_0$ is the scaling constant, $R$ is the universal gas constant and $T$ is the absolute temperature. Here, $\Delta G(T)$ represents the temperature-dependent free energy of unfolding, given by:

$$\Delta G(T) = \Delta G_{T_r} - \delta T \Delta S_{T_r} \tag{S8}$$

where $T_r$ is a reference temperature, $\Delta S_{T_r}$ denotes the entropy of unfolding at $T_r$, $\Delta G_{T_r}$ denotes free energy of unfolding at $T_r$ and $\delta T = T - T_r$. Detailed derivation can be found in Chen and Shakhnovich (2010).

**Background**

Johnson and Lewin (Johnson and Lewin, 1946b) observed that Escherichia coli grown at 45 °C ceased growth but resumed exponential proliferation upon transfer to 37 °C. They proposed that cells undergo reversible damage and hypothesized that a single master enzyme ($E_n$) with free energy of unfolding $\Delta G$ controls the population growth rate:

$$s(T) = E_n \frac{k_B T}{h} e^{-\Delta G/RT} \tag{S9}$$

The enzyme transitions between native and denatured forms in a two-state folding model:

$$E_n \rightleftharpoons^{k_1}_{k_2} E_d \tag{S10}$$

In its native state, a protein exhibits low enthalpy and conformational entropy, both of which increase upon denaturation. Protein stability arises from intra-protein and protein-solvent interactions, which shift during folding/unfolding. A protein unfolds when the $\Delta G$ between the folded and unfolded states becomes zero. Point mutations predominantly affect $\Delta G$ the of the native state (Zeldovich et al., 2007).

The equilibrium ratio of concentrations is given by:

$$\frac{k_1}{k_2} = \frac{E_n}{E_d} = e^{-\Delta G/RT} \tag{S11}$$

Letting $E_0$ denote the total enzyme concentration:

$$E_n = \frac{E_0}{1 + e^{-\Delta G/RT}} \tag{S12}$$

Substituting Eq.S12 into Eq.S9 yields:

$$s(T) = \frac{E_0 k_B T}{h} \cdot \frac{e^{-\Delta G^{\ddagger}/RT}}{1 + e^{-\Delta G/RT}} \tag{S13}$$

Chen and Shakhnovich (2010) extend this model by associating the replication rate of an organism with the functionality of proteins encoded by its essential genes. This assumption is supported by knockout experiments showing that disabling a small number of essential genes can result in lethality (Fraser et al., 2000). Hence, viability is defined by the stability of all essential gene products. Notably, protein hyperstability is assumed to be selectively neutral—greater stability does not necessarily imply higher fitness.

The fractional stability of a rate-determining protein (RDP) with $\Delta G$ is:

$$f_i = \frac{1}{1 + e^{-\Delta G/RT}} \tag{S14}$$

Consequently, the temperature-dependent scaling of growth rate assuming a single RDP becomes:

$$s(p_{\text{evo}} = [\Delta G_{T_r}, \Delta S_{T_r}], T) = \begin{cases} b_0 \dfrac{e^{-\Delta H^{\ddagger}/RT}}{1 + e^{-\Delta G(T)/RT}}, & \text{if } \Delta G(T) < 0 \\ 0, & \text{otherwise} \end{cases} \tag{S15}$$

where $\Delta H^{\ddagger}$ is the metabolic free energy barrier. The temperature dependence of $\Delta G$ is given by:

$$\Delta G(T) = \Delta H_{T_r} + \Delta C_p (T - T_r) - T \Delta S_{T_r} - \Delta C_p T \ln\left(\frac{T}{T_r}\right) \tag{S16}$$

Here, $T_r$ is the reference temperature, $\Delta H_{T_r}$ is the enthalpy of unfolding at $T_r$, $\Delta S_{T_r}$ is the entropy of unfolding at $T_r$ and $\Delta C_p$ is the constant heat capacity difference. This expression can be recast as:

$$\Delta G(T) = \Delta G_{T_r} + \Delta C_p \delta T - \delta T \Delta S_{T_r} + \Delta C_p T_r \ln\left(\frac{T_r + \delta T}{T_r}\right) \tag{S17}$$

where $\delta T = T - T_r$. Given that $\delta T / T_r \ll 1$, we simplify the expression to a linear form:

$$\Delta G(T) = \Delta G_{T_r} - \delta T \Delta S_{T_r} \tag{S18}$$

#### Enzyme catalysis-based model

Several models have attempted to mechanistically explain the unimodal TPCs observed in enzymatic reactions (DeLong et al., 2017; Hobbs et al., 2017; Chen and Shakhnovich, 2009; Johnson and Lewin, 1946c; Schoolfield et al., 1981; Frazier et al., 2006). Although protein denaturation has often been proposed as a key mechanism underlying the decline in enzyme activity at high temperatures, experimental data suggest that catalytic efficiency tends to decline more sharply than expected from thermal instability alone (Somero, 2004; Daniel and Danson, 2010). This is particularly true for cold-adapted enzymes, which may lose function at relatively low temperatures while remaining structurally intact.

To address this, Hobbs et al. (2017) introduced the Macromolecular Rate Theory (MMRT), henceforth referred to as the Enzyme catalysis-based model. Based on transition state theory (Eyring, 1935), MMRT attributes the unimodality of enzyme kinetics not to denaturation, but to the intrinsic thermodynamic properties of enzyme-substrate interactions—specifically, the temperature dependence of the Gibbs free energy of activation.

A key observation from Hobbs et al. (2017) was that many enzymes exhibit negative heat capacity changes during catalysis ($\Delta C_p^\ddagger < 0$), consistent with strong enzyme-ligand binding in the transition state, which is typical of macromolecules. These negative valued $\Delta C_p^\ddagger$ were also found to be largely temperature-independent. This allowed the theory to explain the observed decline in reaction rates at high temperatures, including the curvature of TPCs, without invoking protein denaturation as the limiting factor. This is particularly relevant for cold-adapted enzymes, which often lose activity well before denaturing.

According to Eqn. S2, the catalytic rate constant $k_T$ is given by:

$$k_T = \frac{k_B T}{h} e^{\Delta \frac{S_T^\ddagger}{R}} e^{\Delta \frac{-H_T^\ddagger}{RT}} \tag{S19}$$

where $\Delta G_T^\ddagger = \Delta H_T^\ddagger - T\Delta S_T^\ddagger$ is the Gibbs free energy of activation, $\Delta H_T^\ddagger$ is enthalpy of activation and $\Delta S_T^\ddagger$ is entropy of activation at temperature $T$ respectively. The temperature dependence of these quantities is governed by a constant heat capacity change $\Delta C_p^\ddagger$ following Eqns. **??**:

$$\Delta H^\ddagger(T) = \Delta H_{T_r}^\ddagger + \Delta C_p^\ddagger (T - T_r) \tag{S20}$$

$$\Delta S^\ddagger(T) = \Delta S_{T_r}^\ddagger + \Delta C_p^\ddagger \ln\left(\frac{T}{T_r}\right) \tag{S21}$$

where $T_r$ is the reference temperature. Thus, the temperature-scaling of the population growth rate, assuming a catalysis-limited process, is given by:

$$s(p_{\mathrm{evo}} = [\Delta H^{\ddagger}_{T_r}, \Delta S^{\ddagger}_{T_r}, \Delta C^{\ddagger}_{p}], T) = \begin{cases} \frac{k_B T}{h} \exp\left(\frac{\Delta S^{\ddagger}(T)}{R}\right) \exp\left(\frac{-\Delta H^{\ddagger}(T)}{RT}\right), & \text{if } \Delta S^{\ddagger}(T) < 0 \\ 0, & \text{otherwise} \end{cases} \tag{S22}$$

The Enzyme catalysis-based model has since been applied to characterize microbial growth rates in soil ecosystems (Alster et al., 2016) and leaf respiration in plants (Liang et al., 2018), further supporting its applicability across biological systems where enzyme function limits metabolic performance.

In summary, the Enzyme catalysis-based model offers a thermodynamically grounded explanation for temperature-dependent reaction rates, particularly in cases where enzymatic activity declines precede protein denaturation.

##### Gaussian Model

We also provide a reference or control for the effect of asymmetry in the thermodynamically motivated TPCs defined above. Hence, we compare the mechanistic TPCs with a Gaussian TPC with mean $T_0$ and standard deviation $\sigma$. The temperature scaling function is given by:

$$s(p_{evo} = [T_0, \sigma], T) = b_0 e^{\frac{(T-T_0)^2}{2(\sigma)^2}} \tag{S23}$$

Evolving parameters are the mean $T_0$ and standard deviation $\sigma$.

#### Thermal adaptation

Understanding thermal adaptation in proteins is critical for elucidating macroecological patterns of species distributions. This section explores the two models: the Protein denaturation-based model and the Enzyme catalysis-based model. Both models focus on the interplay between thermodynamic parameters, such as activation free energy, activation entropy, and heat capacity change, under varying thermal environments. By investigating how these parameters evolve in response to selective pressures, we gain insight into the mechanisms by which organisms adapt to extreme temperatures. Empirical evidence highlights the significance of these adaptations in enhancing protein stability and catalytic efficiency, ultimately shaping evolutionary trajectories and ecological dynamics. We will elaborate on each model, examining the specific roles of the parameters involved and their evolutionary implications.

##### Protein denaturation-based model

The Protein denaturation-based model (Eqn. S7) incorporates three key parameters: the free energy for unfolding ($\Delta G_{T_r}$), the entropy of unfolding ($\Delta S_{T_r}$), and the exponential growth rate scaling factor $\Delta H^{\ddagger}$, with all thermodynamic quantities evaluated at a reference temperature $T_r$. Among these, $\Delta G_{T_r}$ and $\Delta S_{T_r}$ are of particular interest for understanding thermal adaptation through protein stability, and their evolutionary modulation is central to shaping macroecological patterns across diverse taxa.

Empirical evidence supports the adaptive evolution of $\Delta G_{T_r}$, which reflects protein thermal stability. Point mutations often shift $\Delta G_{T_r}$, making it a mutable and evolutionarily relevant trait (Zeldovich et al., 2007). Organisms inhabiting high-temperature environments tend to evolve more thermally stable proteins, characterized by elevated $\Delta G_{T_r}$ values. For instance, Stark et al. (2022) analyzed a broad dataset of enzyme catalytic rate constants and found a strong correlation between enzyme melting temperatures and optimal growth temperatures. This suggests that protein thermostability, rather than catalytic compensation, is the predominant mode of thermal adaptation at high temperatures.

In contrast, cold adaptation often involves changes in $\Delta S_{T_r}$, which modulates the entropy associated with protein folding. Entropy-increasing mutations promote structural flexibility, a key requirement for maintaining enzymatic function at low temperatures. Studies on psychrophilic enzymes support this pattern: mutations that elevate entropy facilitate catalysis under cold conditions (Fields and Somero, 1998; Siddiqui, 2017). Similarly, in Escherichia coli, entropy-modulating mutations in adenylate kinase were shown to improve function in cold environments (Saavedra et al., 2018).

Taken together, these findings highlight the dual roles of $\Delta G_{T_r}$ and $\Delta S_{T_r}$ in thermal adaptation. While increased stability (higher $\Delta G_{T_r}$) underlies adaptation to heat, enhanced flexibility (via changes in $\Delta S_{T_r}$) facilitates cold tolerance. These complementary strategies illustrate how organisms fine-tune protein energetics to match their thermal niches.

Accordingly, we treat both $\Delta G_{T_r}$ and $\Delta S_{T_r}$ as evolving traits under selection in our simulations. The Gaussian mutation kernels and initial standing variation for these parameters are provided in Table S2.

##### Enzyme catalysis-based model

The Enzyme catalysis-based model (Hobbs et al., 2017), described by Eqn. S22, incorporates three key parameters: the activation enthalpy ($\Delta H^{\ddagger}_{T_r}$), activation entropy ($\Delta S^{\ddagger}_{T_r}$) at a reference temperature $T_r$, and the heat capacity change ($\Delta C^{\ddagger}_p$). Together, these parameters govern the Gibbs free energy of activation and thus determine enzymatic performance across temperatures. Evolutionary changes in these traits can enable organisms to optimize enzyme activity for different thermal environments.

In cold-adapted (psychrophilic) enzymes, for example, lower activation enthalpy promotes higher catalytic turnover at low temperatures (Low et al., 1973; Fields and Somero, 1998). Thermophilic enzymes, on the other hand, often exhibit higher activation enthalpy, enhancing structural stability at elevated temperatures. Phosphoglucose isomerase (Pgi), an essential enzyme in glycolysis and gluconeogenesis, illustrates this pattern. Cold-adapted Pgi variants typically display lower activation enthalpy and higher entropy, increasing catalytic efficiency under cold conditions, while heat-adapted variants show the opposite trend—reflecting a trade-off between kinetic efficiency and thermostability (Bomblies and Peichel, 2022).

This trade-off is further governed by enthalpy–entropy compensation (EEC), a well-documented phenomenon where changes in binding strength and flexibility offset one another thermodynamically (Rosenberg et al., 1971; Liu and Guo, 2001; Fox et al., 2018). Stronger enzyme-substrate binding often lowers enthalpy but reduces conformational entropy, while increased flexibility may raise entropy at the cost of binding affinity. EEC manifests as a linear relationship between $\Delta H^{\ddagger}_{T_r}$ and $\Delta S^{\ddagger}_{T_r}$, first observed in catalytic systems by Constable (1925) and further analyzed in protein systems (Rosenberg et al., 1971; Fox et al., 2018; Dragan et al., 2017).

To reflect this, we assume a linear constraint between $\Delta H^{\ddagger}_{T_r}$ and $\Delta S^{\ddagger}_{T_r}$ in our model:

$$\Delta S^{\ddagger}_{T_r} = \frac{(\Delta H^{\ddagger}_{T_r} - 0.9)}{330} \tag{S24}$$

This assumption follows earlier applications, such as in the Hinshelwood model fitted to phytoplankton TPCs by Grimaud et al. (2017).

The third parameter, $\Delta C^{\ddagger}_{p}$, governs the curvature of the TPC and reflects the temperature-dependence of both $\Delta H^{\ddagger}$ and $\Delta S^{\ddagger}$. Increasing $\Delta C^{\ddagger}_{p}$ can shift the temperature optimum upward, which is commonly observed in both inter- and intraspecific comparisons of thermal adaptation (Somero, 1995; Hochachka and Somero, 2002; Schulte et al., 2015; Bomblies and Peichel, 2022). For instance, Hobbs et al. (2017) showed that targeted mutagenesis of barnase, as well as comparative data from IPMDH enzymes in Bacillus species, illustrated such thermoadaptive shifts.

Additionally, evolutionary reconstructions by Nguyen et al. (2017) on adenylate kinase (Adk) across 3 billion years revealed that adaptation to a cooling Earth primarily occurred through reductions in $\Delta C^{\ddagger}_{p}$, rather than through enthalpy minimization. This reduced the temperature sensitivity of catalytic rates, allowing enzymes to remain functional under lower thermal regimes.

Given this empirical basis, we allow all three MMRT parameters—$\Delta H^{\ddagger}_{T_r}$, $\Delta S^{\ddagger}_{T_r}$, and $\Delta C^{\ddagger}_{p}$—to evolve in our simulations. Their mutation kernels and initial standing variation are summarized in Table S2.

#### Individual-based model

Following the modified Gillespie algorithm by Allen and Dytham, 2009 (Allen and Dytham, 2009), we simulate Eq. 5. In each time step an individual may experience one of the three events: birth, death or emigration. An individual's traits define its birth probability ($b_i$), death probability ($d_i$) and emigration ($e_i$).

To understand the modification we first describe a direct Gillespie algorithm:

1. Probabilities of giving birth, death or dispersing for an individual $i$ are given by its trait $b_i, d_i, e_i$, respectively.

2. Each time-step is picked from an exponential distribution with mean $\lambda = \Sigma(b_i + d_i + e_i)$.

3. The next event is calculated where The probability of birth, death or dispersal for individual $j$ is $\frac{b_j}{\Sigma(b_i+d_i+e_i)}$, $\frac{d_j}{\Sigma(b_i+d_i+e_i)}$ and $\frac{d_j}{\Sigma(b_i+d_i+e_i)}$ respectively.

Each event requires rates to be calculated for every member of the population. In the modified Gillespie algorithm, per-event calculation is independent of population size, which implies an important gain in simulation time:

1. Probabilities of giving birth, death or dispersing for an individual $i$ are given by its trait $b_i, d_i, e_i$, respectively.

2. Each time-step is picked from an exponential distribution with mean $\lambda = (c_b + c_d + c_e)N$ where $c_b >= max(b_i : \forall i)$, $c_e >= max(e_i : \forall i)$ and $c_d >= max(d_i : \forall i)$.

3. The event is selected where The probability of birth, death or dispersal is $\frac{c_b}{(c_b+c_e+c_d)}$, $\frac{c_e}{(c_b+c_e+c_d)}$ and $\frac{c_d}{(c_b+c_e+c_d)}$ respectively.

4. For individual $j$ the The probability of the selected event occurring is $\frac{b_j}{c_b}$, $\frac{e_j}{c_e}$ and $\frac{d_j}{c_d}$ respectively.

5. The next event is selected where the probability of birth, death or dispersal for individual $j$ is $\frac{c_b}{(c_b+c_e+c_d)}$, $\frac{c_e}{(c_b+c_e+c_d)}$ and $\frac{c_d}{(c_b+c_e+c_d)}$, respectively.

##### Birth rate

An event is selected by relative maximum rate constants. Hence, the probability of birth event is :

$$\frac{c_b}{(c_b + c_d + c_e)} \tag{S25}$$

taking density regulation into account, The probability of the birth event being executed is given by:

$$\frac{b_i}{(1+\beta_i N)c_b} \tag{S26}$$

**Death rate**

All individuals have equal death rates. The probability of choosing a death event is given by:

$$\frac{c_d}{(c_b+c_d+c_e)} \tag{S27}$$

The probability of executing a death event is given by:

$$\frac{d_i}{c_d} \tag{S28}$$

**Emigration rate**

Emigration is density-independent. The probability of choosing an emigration event is given by:

$$\frac{c_e}{(c_b+c_d+c_e)} \tag{S29}$$

The probability of executing an emigration event is given by:

$$\frac{e_i}{c_e} \tag{S30}$$

A dispersal cost is applied as emigration mortality $e_\mu$ if executed.

### Supplementary Figures

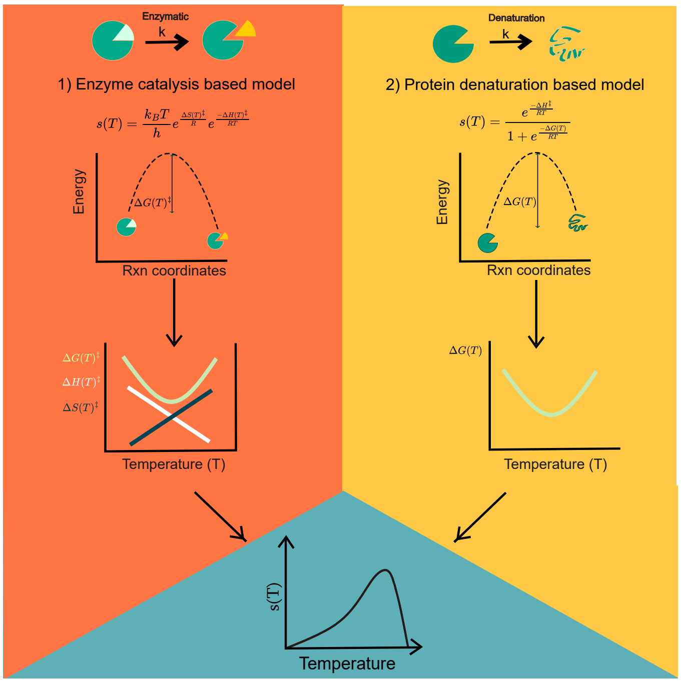


Figure S1: The Eyring-Evans-Polanyi (EEP) equations (Eyring, 1935), derived from Transition State Theory, describe the formation of a high-energy transition state during chemical reactions. These equations use the temperature dependence of enthalpy and entropy changes (Eqns. S6 and S5) to explain how reaction rates vary with temperature. We explore two models from the literature that apply the EEP equations to explain the unimodality of temperature performance curves (TPCs) for population growth rates ($b_0 s(T)$): 1) Protein denaturation-based model by Chen and Shakhnovich (2010): This model uses the EEP equations to describe the temperature dependence of protein denaturation in terms of Gibbs free energy ($\Delta G$), enthalpy change ($\Delta H$), and entropy change ($\Delta S$). It estimates the fraction of a rate-determining protein in its folded state as $1/(1 + e^{\frac{-\Delta G(T)}{RT}})$, which directly controls metabolic activity rates ($e^{\frac{-\Delta H^\ddagger}{RT}}$). 2) Enzyme catalysis-based model by Arcus et al. (2016): This model employs the EEP equations to describe the temperature dependence of enzyme catalysis reactions, using Gibbs free energy of activation ($\Delta G^\ddagger$), enthalpy change of activation ($\Delta H^\ddagger$), and entropy change of activation ($\Delta S^\ddagger$).

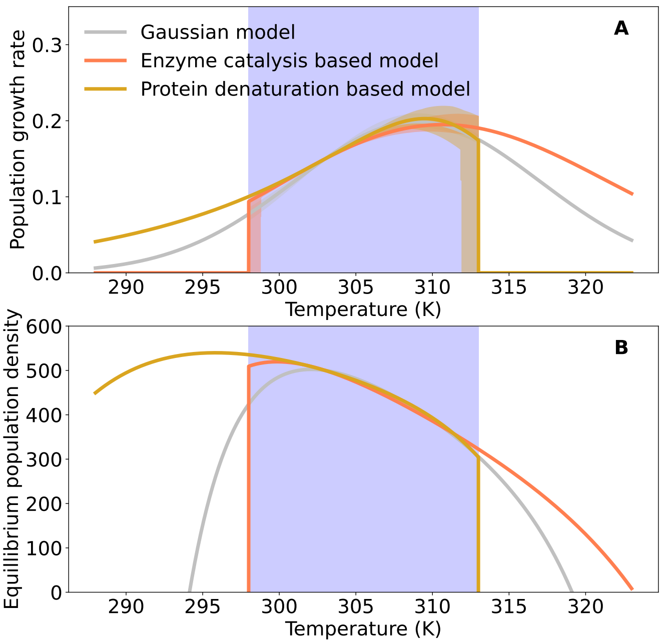


Figure S2: A) Initial thermal performance curves for the three models after fitting Eqn. S7 for the Protein denaturation-based model and Eqn. S22 for the Enzyme catalysis-based model to Eqn. S23 for the Gaussian model in the shaded thermal range. B) Corresponding equilibrium density given by Eqn. 6 across the landscape, neglecting the effect of dispersal and edge effects. Population growth parameter values can be found in Table S1. Model-wise temperature scaling parameters can be found in Table S2.

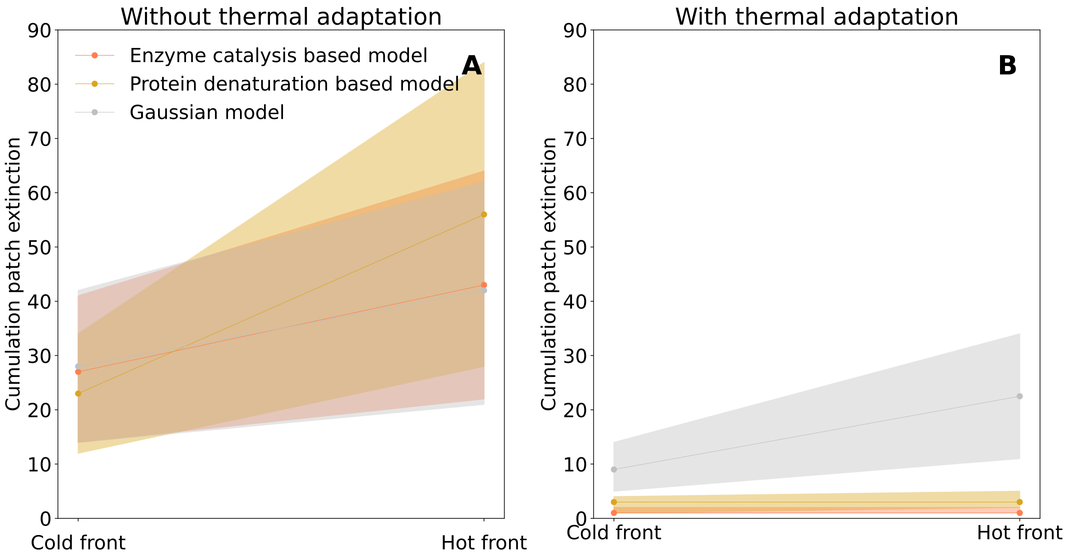


Figure S3: Patch extinctions at the hot and cold patch front during range expansions. A) Cumulative patch extinctions at cold and hot front in simulations with thermal adaptation. B) Cumulative patch extinctions at cold and hot front in simulations without thermal adaptation. The number of extinctions at the front are added up across the duration of the simulation. The shaded region gives the interquartile range of the number of extinctions among replicates.

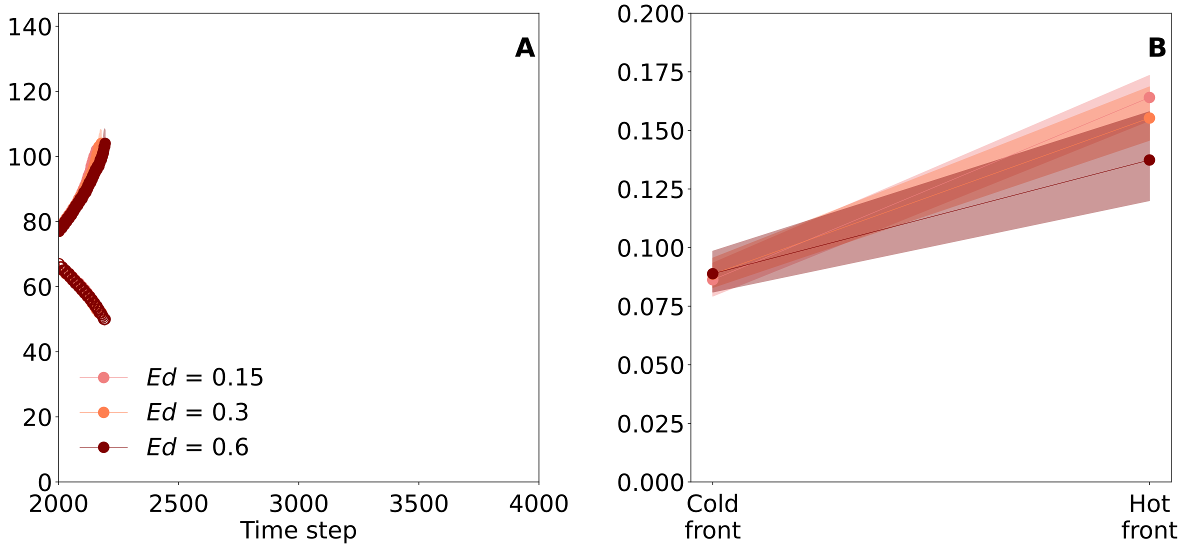


Figure S4: Sensitivity analysis of the Enzyme catalysis-based model to higher death rate coefficient with $E_d = 0.6$ and to lower death rate coefficient $E_d = 0.15$. Patch fronts are averaged across 80 replicates. At every 2 time steps, the last 3 occupied patches with more than 10 individuals are considered as patch front. A): Range front position versus time point. The shaded region gives the inter-quartile range of the front among replicates; B) : Initial speeds of expansion at the cold and hot patch front calculated by fitting a linear curve to patch front dynamics before 2100 time steps. Population growth parameter values can be found in Table S1. Model wise temperature scaling parameters can be found in in Table S2.

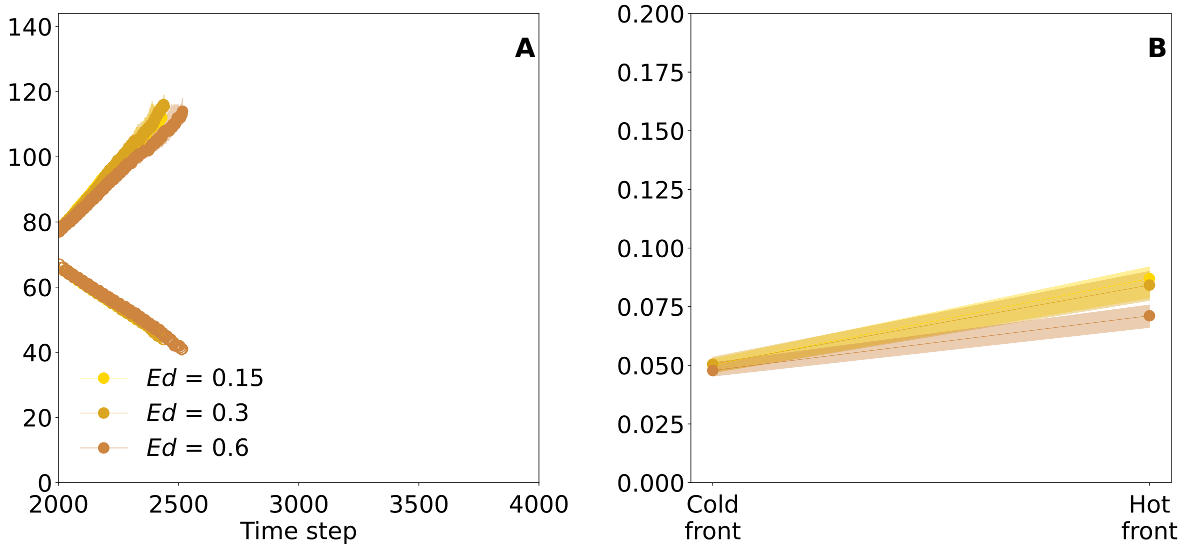


Figure S5: Sensitivity analysis of the Protein denaturation-based model to higher death rate coefficient with $E_d = 0.6$ and to lower death rate coefficient $E_d = 0.15$. Patch fronts are averaged across 80 replicates. At every 2 time steps, the last 3 occupied patches with more than 10 individuals are considered as patch front. A): Range front position versus time point. The shaded region gives the inter-quartile range of the front among replicates; B) : Initial speeds of expansion at the cold and hot patch front calculated by fitting a linear curve to patch front dynamics before 2100 time steps. Population growth parameter values can be found in Table S1. Model wise temperature scaling parameters can be found in in Table S2.

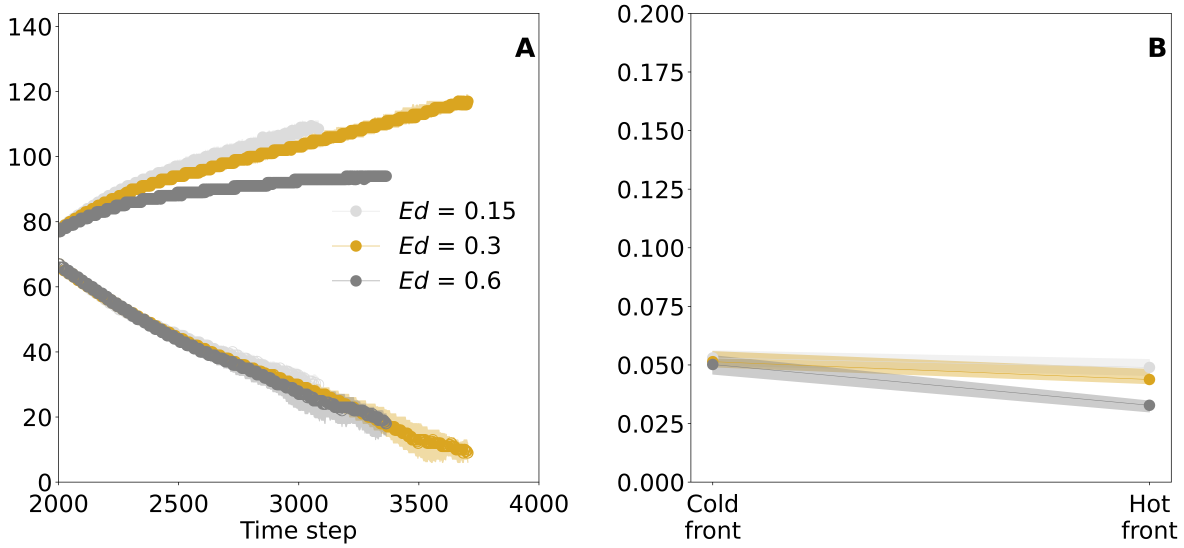


Figure S6: Sensitivity analysis of the Gaussian model to a higher death rate coefficient with $E_d = 1.2$ and to lower death rate coefficient $E_d = 0.3$. Patch fronts are averaged across 80 replicates. At every 2 time steps, the last 3 occupied patches with more than 10 individuals are considered as patch front. A): Range front position versus time point. The shaded region gives the inter-quartile range of the front among replicates; B) : Initial speeds of expansion at the cold and hot patch front calculated by fitting a linear curve to patch front dynamics before 2100 time steps. Population growth parameter values can be found in Table S1. Model wise temperature scaling parameters can be found in in Table S2.

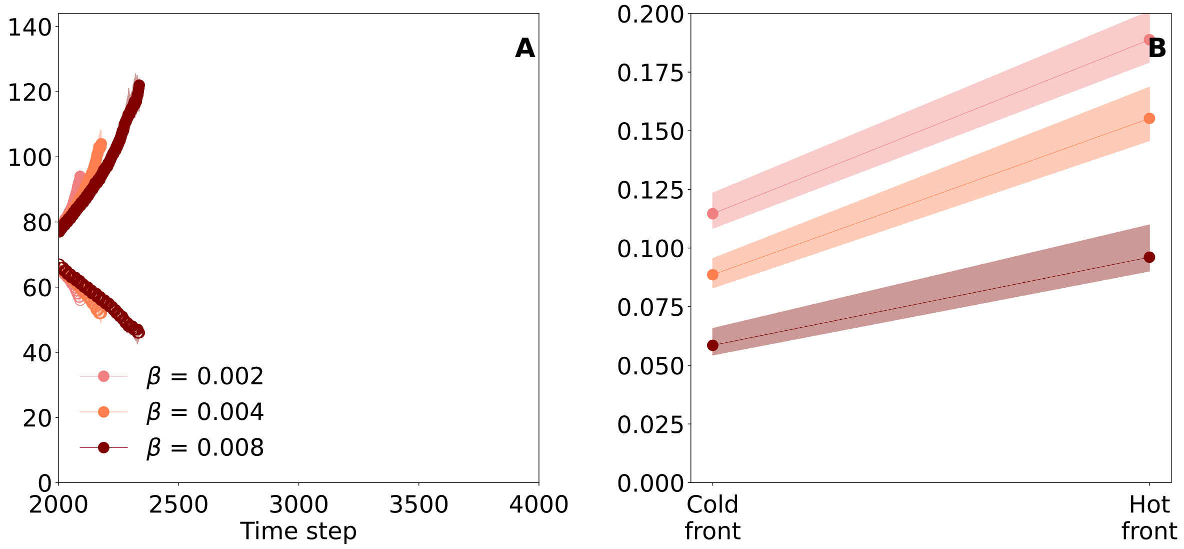


Figure S7: Sensitivity analysis of the Enzyme catalysis-based model to higher intraspecific competition coefficient with $\beta = 0.008$ and to lower intraspecific competition coefficient $\beta = 0.002$. Patch fronts are averaged across 80 replicates. At every 2 time steps, the last 3 occupied patches with more than 10 individuals are considered as patch front. A): Range front position versus time point. The shaded region gives the inter-quartile range of the front among replicates; B) : Initial speeds of expansion at the cold and hot patch front calculated by fitting a linear curve to patch front dynamics before 2100 time steps. Population growth parameter values can be found in Table S1. Model wise temperature scaling parameters can be found in in Table S2.

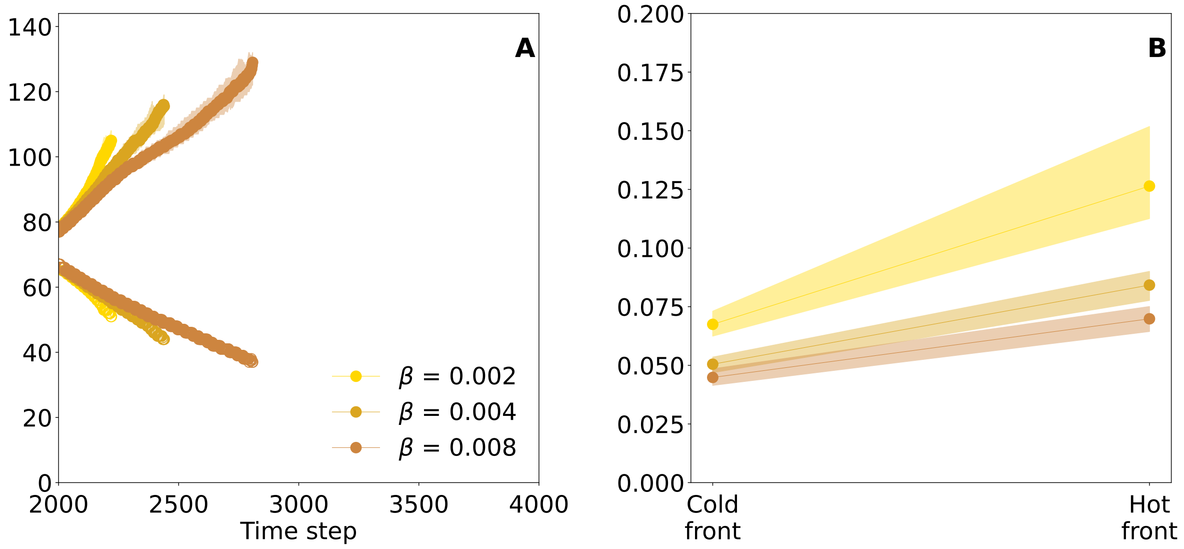


Figure S8: Sensitivity analysis of the Protein denaturation-based model to higher intraspecific competition coefficient with $\beta = 0.008$ and to lower intraspecific competition coefficient $\beta = 0.002$. Patch fronts are averaged across 80 replicates. At every 2 time steps, the last 3 occupied patches with more than 10 individuals are considered as patch front. A): Range front position versus time point. The shaded region gives the inter-quartile range of the front among replicates; B) : Initial speeds of expansion at the cold and hot patch front calculated by fitting a linear curve to patch front dynamics before 2100 time steps. Population growth parameter values can be found in Table S1. Model wise temperature scaling parameters can be found in in Table S2.

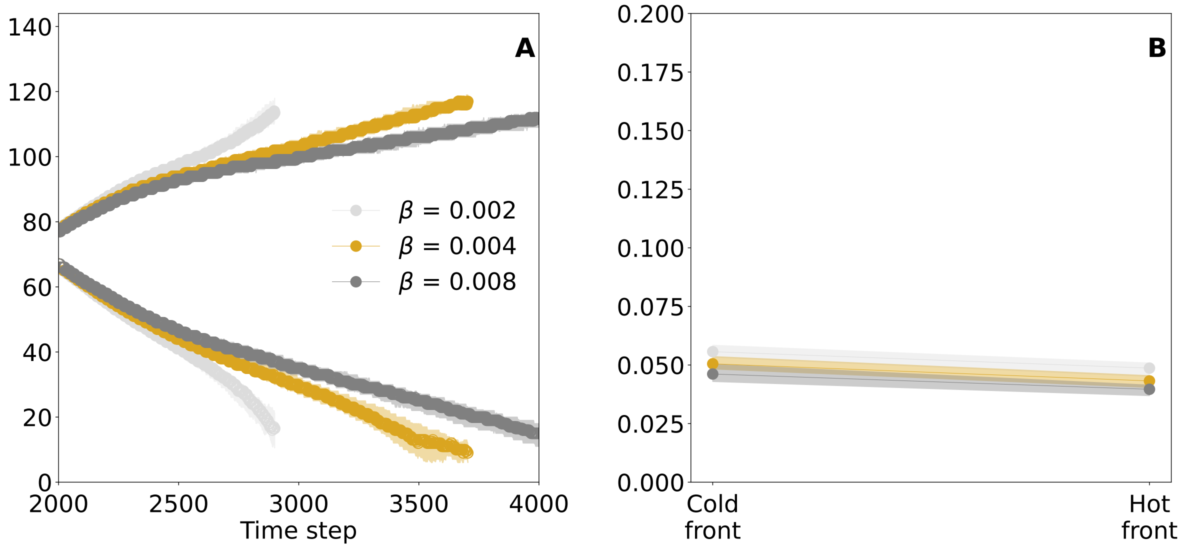


Figure S9: Sensitivity analysis of the Gaussian model to higher intraspecific competition coefficient with $\beta = 0.008$ and to lower intraspecific competition coefficient $\beta = 0.002$. Patch fronts are averaged across 80 replicates. At every 2 time steps, the last 3 occupied patches with more than 10 individuals are considered as patch front. A): Range front position versus time point. The shaded region gives the inter-quartile range of the front among replicates; B) : Initial speeds of expansion at the cold and hot patch front calculated by fitting a linear curve to patch front dynamics before 2100 time steps. Population growth parameter values can be found in Table S1. Model wise temperature scaling parameters can be found in in Table S2.

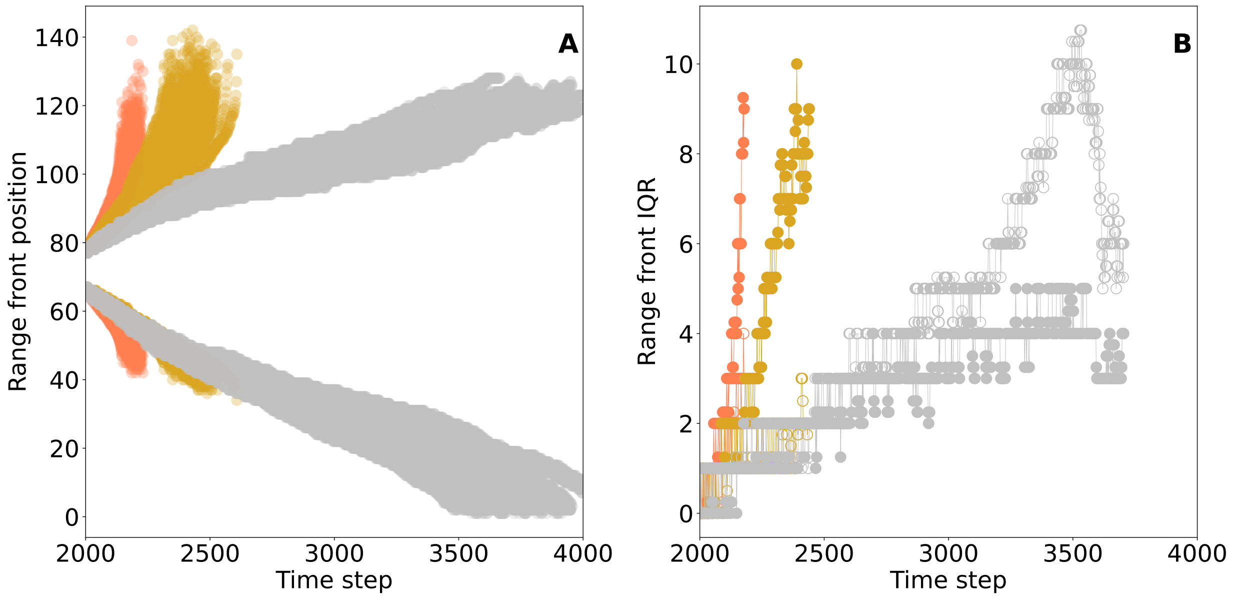


Figure S10: Variation in range expansion trajectories. A) Range expansion patch front position for each of the 80 replicates with median line graph overlaid. The shaded region gives the interquartile range among replicates. B) Interquartile range of the range front position for each model plotted with respect to time. Unfilled markers belong to cold front expansions and filled markers belong to hot front expansion. Population growth parameter values can be found in Table S1. Model wise temperature scaling parameters can be found in in Table S2.

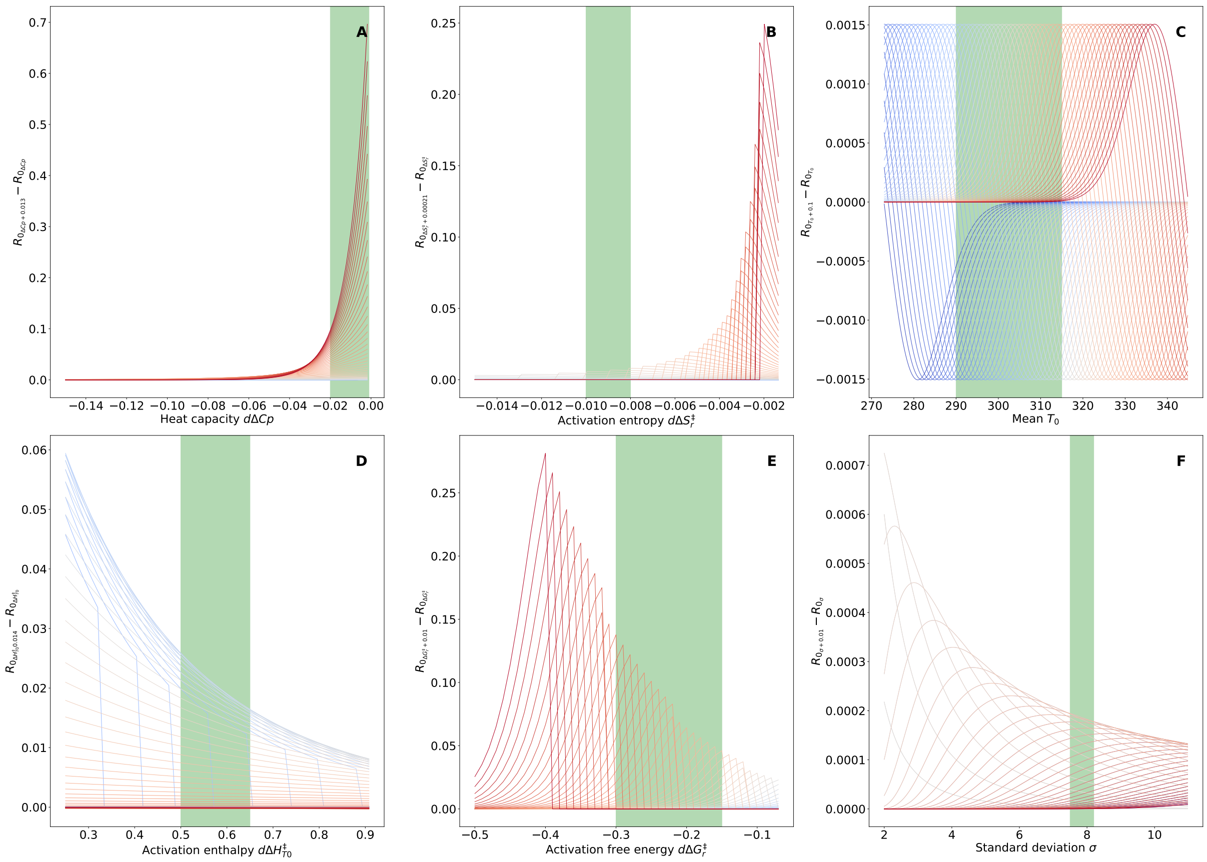


Figure S11: Numerical properties of all three models. For each graph, on the y-axis we have the change in growth rate for a small change in the evolvable trait. The x-axis shows the evolvable trait. Fig. A and B - Each curve is numerically plotted for the Enzyme catalysis-based model along a gradient of temperatures represented by colors of the graphs. A): Temperature dependence of mutation effects for Heat capacity change $\Delta C_p$ ; B): Temperature dependence of mutation effects for Activation enthalpy, $\Delta H^{\ddagger}_{T_r}$. Fig C and D - Each curve is numerically plotted for the Protein denaturation-based model along a gradient of temperatures represented by colors of the graphs. A): Temperature dependence of mutation effects for Activation free energy, $\Delta G_{T_r}$ ; B): Temperature dependence of mutation effects for Activation entropy, $\Delta S_{T_r}$. Fig E and F - Each curve is numerically plotted for the Protein denaturation-based model along a gradient of temperatures represented by colors of the graphs. E): Temperature dependence of mutation effects for Mean temperature, $T_0$ ; F): Temperature dependence of mutation effects for Mean temperature, $T_0$. The shaded green rectangle shows a range of the parameter value achieved in our simulations. Population growth parameter values can be found in Table S1. Model wise temperature scaling parameters can be found in in Table S2.

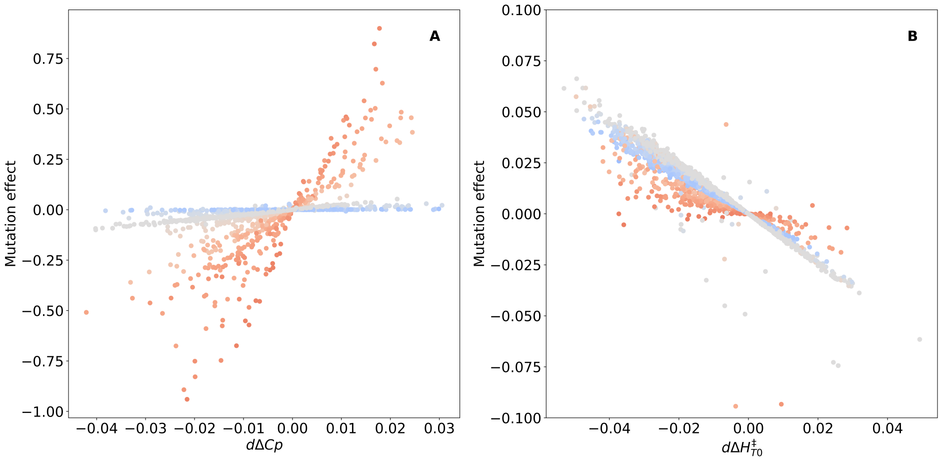


Figure S12: We show the individual contribution of mutations in evolving trait to changed in growth rate. The y-axis has the mutation effect of the change in growth rate with respect to a mutation in the trait, whose magnitude is shown on the x-axis. The color of the points denotes the temperature of the patch where the mutation occurs. They are in accordance with Fig. S11 A): Mutation effect versus mutation in Heat capacity change $\Delta C_p$ ; B): Mutation effect versus mutation in Activation entropy, $\Delta S_{T_r}$.

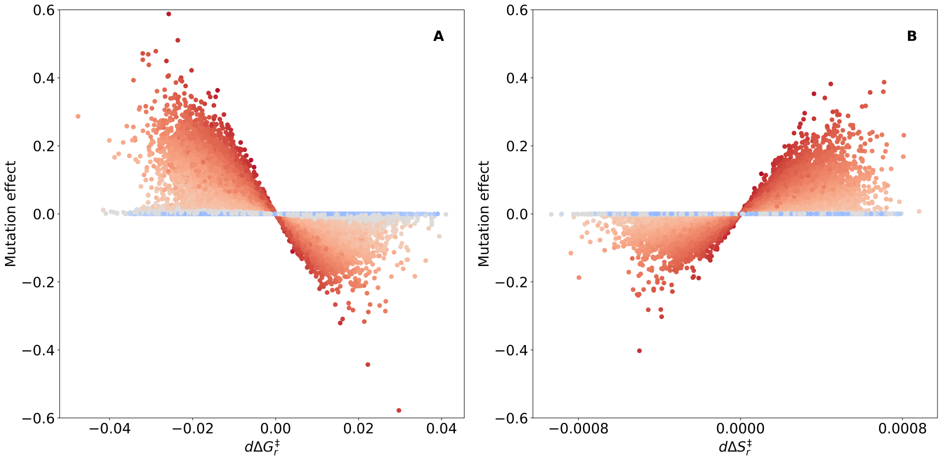


Figure S13: We show individual contribution of mutations in evolving trait to changed in growth rate. The y-axis has the mutation effect of the change in growth rate with respect to a mutation in the trait whose magnitude is shown on the x axis. The color of the points denotes the temperature of the patch where the mutation occurs. They are in accordance with Fig. S11 A): Mutation effect versus mutation in free energy of unfolding, $\Delta G_{T_r}$ ; B): Mutation effect versus mutation in entropy of unfolding, $\Delta S_{T_r}$.

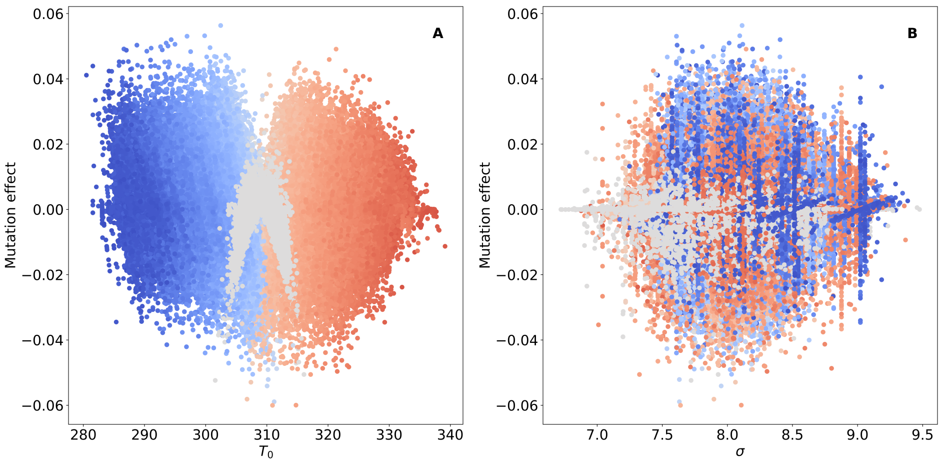


Figure S14: We show individual contribution of mutations in evolving trait to changed in growth rate. The y-axis has the mutation effect of the change in growth rate with respect to a mutation in the trait whose magnitude is shown on the x axis. The color of the points denotes the temperature of the patch where the mutation occurs. They are in accordance with Fig. S11 A): Mutation effect versus mutation in Mean temperature, $T_0$ ; B): Mutation effect versus mutation in Mean temperature, $T_0$.

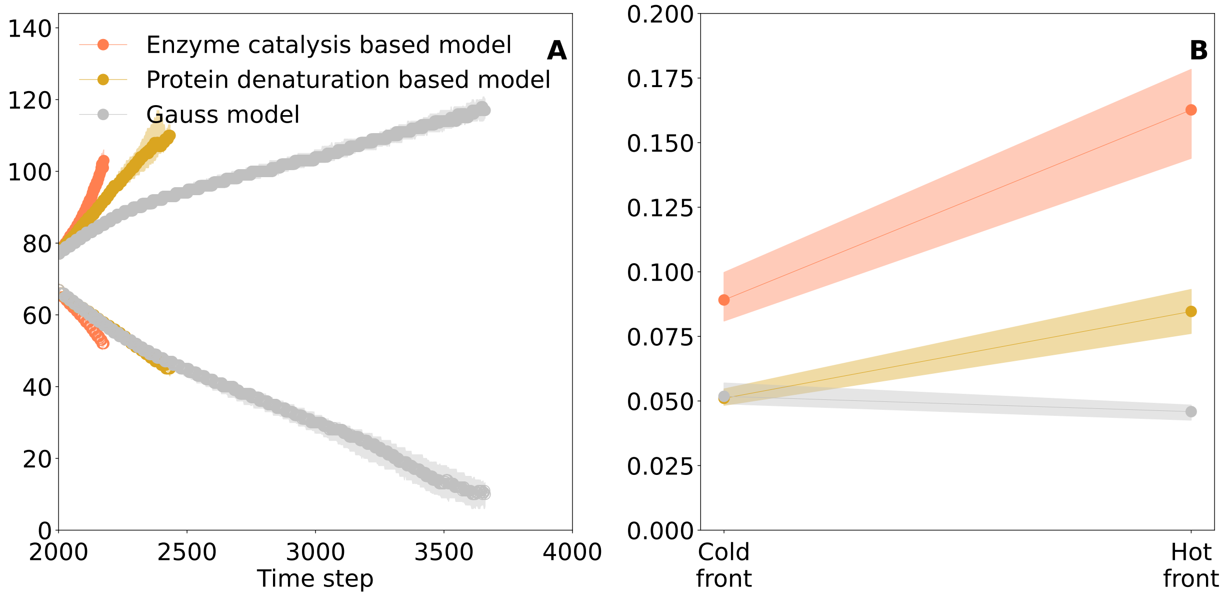


Figure S15: Population dynamics from simulations with thermal adaptation and dispersal evolution. Patch fronts are averaged across 80 replicates. Every 2 time steps, the last 3 occupied patches with more than 10 individuals are considered patch front. A): Population density versus time steps. The shaded region gives the interquartile range among replicates; B): Initial speeds of expansion at the cold and hot patch front calculated by fitting a linear curve to patch front dynamics before 2100 time steps. Population growth parameter values can be found in Table S1. Model wise temperature scaling parameters can be found in in Table S2.

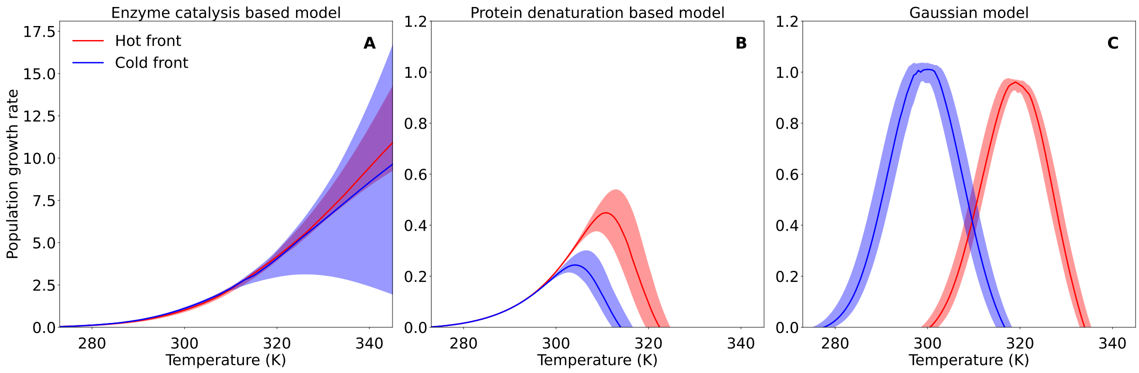


Figure S16: Comparison of TPC at patch fronts with dispersal evolution. Hot front, cold front and core patch's niches was calculated for individuals at the patch front. A) TPC of Enzyme catalysis-based model, B) TPC of Protein denaturation-based model, C) TPC of Gaussian model. Shaded area is the variance of the thermal niche between the 80 replicates.

#### Supplementary Tables

Table S1: Common parameters for all simulations. Emigration rate is utilized for the dispersal trait when it is not mutating. The mutation kernel is assumed to be a Gaussian distribution with a mutation probability of 0.025.

| IBM Parameters | Description | Value |
|---|---|---|
| $N_x$ | Population density in patch $x$ | Initial value: 200 for patch 67-77 |
| $\beta_0 s(p_{evo}, T_{ref})$ | Intraspecific competition coefficient at $T_{ref} = 20\,°\mathrm{C}$ | 0.004 |
| $d_0 e^{-\frac{E}{RT_{ref}}}$ | Death rate at $T_{ref} = 20\,°\mathrm{C}$ | 0.05 |
| $b_0 s(p_{evo}, T_{ref})$ | Birth rate at $T_{ref} = 20\,°\mathrm{C}$ | 0.15 |
| $e$ | Emigration rate | 0.01 |
| $e_\mu$ | Dispersal cost | 0.01 |
| $T$ | Temperature | 0-72 °C with 0.5 °C increase per patch |

Table S2: Parameters for the Enzyme catalysis, Protein denaturation, and Gaussian curve models of the Thermal Performance Curve (TPC). All mutational kernels and initialisation are Gaussian distributions with the specified standard deviations (SD). The probability of mutation is 0.05 for each parameter.

| **Protein denaturation-based model** | | |
|---|---|---|
| Parameter | Description | Value |
| $\Delta H^{\ddagger}$ | Metabolic free energy barrier | 0.67 eV |
| $\Delta G_{T_r}$ | Gibbs free energy at 20 °C | 0.091 eV |
| $\Delta G_{T_r}^{mut}$ | Mutation kernel SD of $\Delta G_{T_r}$ | 0.01 eV |
| $\Delta G_{T_r}^{ini}$ | Initial SD of $\Delta G_{T_r} distribution$ | 0.01 eV |
| $\Delta S_{T_r}$ | Activation entropy at 20 °C | 0.0091 eV |
| $\Delta S_{T_r}^{mut}$ | Mutation kernel SD of $\Delta S_{T_r}$ | 0.00021 eV |
| $\Delta S_{T_r}^{ini}$ | Initial SD of $\Delta S_{T_r} distribution$ | 0.00021 eV |
| $E$ | Activation energy for death rate | 0.3 eV, does not evolve |
| **Enzyme catalysis-based model** | | |
| Parameter | Description | Value |
| $\Delta H_{T_r}^{\ddagger}$ | Enthalpy change at 20 °C | 0.74 eV |
| $\Delta H_{T_r}^{mut}$ | Mutation kernel SD of $\Delta H_{T_r}^{\ddagger}$ | 0.014 eV |
| $\Delta H_{T_r}^{ini}$ | Initial SD of $\Delta H_{T_r}^{\ddagger}$ distribution | 0.014 eV |
| $\Delta S_{T_r}^{\ddagger}$ | Entropy change at 20 °C | -0.00054 eV, correlated with $\Delta H_{T_r}^{\ddagger}$ according to Eqn. S24 |
| $\Delta C_p$ | Heat capacity change | -0.072 eV |
| $\Delta C_p^{mut}$ | Mutation kernel SD of $\Delta C_p$ | 0.013 eV |
| $\Delta C_p^{ini}$ | Initial SD of $\Delta C_p$ distribution | 0.013 eV |
| $E$ | Activation energy for death rate | 0.3 eV, does not evolve |
| **Gaussian Curve Model** | | |
| Parameter | Description | Value |
| $T_0$ | Mean of the Gaussian curve | 309 K |
| $T_0^{mut}$ | Mutation kernel SD of $T_0$ | 1 K |
| $T_0^{ini}$ | Initial SD of $T_0$ distribution | 1 K |
| $\sigma$ | SD of the Gaussian curve | 8 K |
| $:\sigma^{mut}$ | Mutation kernel SD of $\sigma$ | 0.1 K |
| $\sigma^{ini}$ | Initial SD of $\sigma$ distribution | 0.1 K |
| $E$ | Activation energy for death rate | 0.3 eV, does not evolve |
| **Dispersal evolution kernel** | | |
| Parameter | Description | Value |
| $e^{mut}$ | Mutation kernel SD of $e$ | 0.001 |
| $e^{ini}$ | Initial SD of $e$ distribution | 0.001 |
